# REVEAL: RNA Ensemble Validation and Exploration with AI-guided Landscapes

**DOI:** 10.64898/2026.11.10.765771

**Authors:** Soumendranath Bhakat, Shray Vats

## Abstract

Capturing the sparsely populated but functionally important, excited states of RNA remains a central challenge for molecular simulation, because the underlying transitions are rare events that unbiased molecular dynamics cannot reach from a single starting structure. Here we introduce REVEAL (RNA Ensemble Validation and Exploration with AI-guided Landscapes), a workflow that couples generative secondary structure sampling, secondary structure conditioned tertiary structure prediction with trRosettaRNA2, unbiased molecular dynamics and Markov state modeling. Using only the sequence of the engineered HIV-1 TAR mutant, in which the excited state ES2 is the dominant conformation, we show that REVEAL captures the reversible ES2 to GS register shift from 6.8 µs of aggregate unbiased sampling across 34 seeds, recovering both basins, the intermediate states, and correctly identifies ES2 as the dominant state, consistent with the design of the construct. In contrast, molecular dynamics seeded from the ES2-dominant HIV-1 TAR mutant NMR ensemble remains trapped in ES2 despite substantially longer aggregate simulation time, and sequence-only ensemble predictors (AlphaFold3, Boltz-2, ESMFold2, OpenFold3 and Protenix-v2) place every structure near the ground state and invert the experimentally observed population ordering. The information content of the seed ensemble, rather than the total simulation time, determines whether the transition is sampled. We further introduce “*population-weighted consensus score*” (PWCS) that ranks conformers by how representative they are of the equilibrium ensemble directly from the simulation data, without experimental restraints and independent of the choice of collective variables. PWCS identifies the dominant state even when low-dimensional representations fail to separate the states. Built entirely from open-source tools, REVEAL provides a transparent and extensible route to RNA conformational ensembles beyond the single-structure paradigm.

## Introduction

RNA molecules play a pivotal role in molecular recognition by transiently storing genetic information, modulating protein synthesis, and regulating gene expression. Unlike globular proteins, the metastable states of RNA molecules are often equally populated and interconvert rapidly. Capturing these transiently populated states is crucial for discovering state-specific RNA therapeutics (small molecules or biologics) beyond the globally optimized single-state paradigm, and for understanding how alternate RNA states regulate molecular recognition^1–4^.

Conformational heterogeneity in RNA can be captured at both the secondary and tertiary structure levels, as the diversity of RNA secondary structures manifests in tertiary structure predictions. A typical strategy for capturing the tertiary structure conformational ensemble is to generate a diverse set of secondary structures and predict the corresponding tertiary structures from each, yielding the final ensemble. De novo RNA tertiary structure prediction models, such as trRosettaRNA2^5^ and FARFAR2^6^, can take secondary structures as input and produce an ensemble of tertiary structures conditioned on the secondary structures. This approach is analogous to MSA subsampling or diffusion/flow matching-based pipelines for globular proteins, in which a conformational ensemble is generated from a given sequence^7^.

Recently, RNAnneal^8^ extended this concept by generating a tertiary structural ensemble of RNA conditioned on its secondary structures and using the ensemble as seeds to initiate molecular dynamics (MD) simulations that capture RNA conformational heterogeneity. In essence, this approach is be inspired by recent methods that launch MD simulations from an initial structural ensemble generated by MSA subsampling^9,10^, diffusion models^11^, or flow matching^12^, such as AlphaFold2-RAVE^13,14^, AlphaFold-MSM^15^, BioEmu-MSM^16^, AlphaFold-SFA^17^, which accelerate the sampling of conformational heterogeneity compared with traditional unbiased MD.

In this work, we introduce a simple and intuitive pipeline that combines a generative AI based RNA tertiary structure predictor, conditioned on the secondary structural ensemble, to generate seeds for unbiased MD simulations. Markov state model^18^ developed using the combined simulation data capture conformational heterogeneity, relative populations, and transition times in RNA. Additionally, we introduce the “*population-weighted consensus score*” (PWCS), which enables the identification of experimentally observed states and of representative macrostates from a pool of metastable states generated by MD simulations. This information can be used in downstream drug discovery workflows, such as virtual screening, de novo protein design, or experimental validation.

The HIV-1 transactivation response element (TAR) is a highly conserved stem-loop that forms at the 5’ end of every nascent viral transcript and is essential for transcriptional elongation from the integrated provirus. The construct used in most biophysical and computational studies comprises of nucleotides 17-45, a 29-nucleotide hairpin built from two A-form helical stems separated by a UCU trinucleotide bulge and capped by a CUGGGA hexanucleotide apical loop^19,20^. The two motifs make up distinct recognition centers: the bulge along with flanking stems engages the arginine rich motif of the viral Tat protein, while the apical loop, together with the bulge, is read out by Cyclin T1 and the super elongation complex (SEC), allowing for P-TEFb dependent elongation^21,22^. As such, HIV-1 TAR transitions between various conformational states to function. The molecule exists in a dynamic equilibrium between a dominant ground state (GS) and two sparsely populated excited states (ES1 and ES2) detectable only by relaxation dispersion NMR. ES1 (∼13% populated, lifetime *∼45* μ*s*) involves a local rearrangement in the apical loop^23^, whereas ES2 (∼0.4% populated, lifetime *∼2 ms*) is a strand shift where the bulge pairs with the upper stem, reshuffling base pairs from the bulge to the tip of the apical loop resulting in three non-canonical base pair mismatches^24,25^. This rearrangement eliminates the motifs recognized by Tat and the SEC, rendering ES2 transcriptionally inactive, and thus, it has been proposed as an auto-inhibited target for conformation-selective therapeutics^26,27^. To allow for this almost invisible state to be studied structurally and computationally, a 31-nucleotide mutant construct (TAR^Mut^) was engineered such that the ES2 fold became the dominant conformation while the GS fold became sparsely populated. Consequently, the transition from ES2 to GS in TAR^Mut^ qualifies as a rare event, expected to lie beyond the sampling capabilities of unbiased molecular dynamics, although its exchange kinetics have not, to our knowledge, been measured directly for this construct.

We used HIV-1 TAR as a model system to investigate RNA conformational heterogeneity, a topic that has been studied extensively with integrated computational and experimental methods. In prior work, a conformational library of RNA tertiary structures was generated using FARFAR2 from the NMR-derived secondary structure^28,29^. Residual dipolar coupling (RDC) data from NMR were then used to select from this library^25^, the conformational ensembles that best matched the experimental data. Enhanced sampling workflows have recently been developed to sample the ES2 ↔ GS transition in TAR^Mut^^30^.

In this work, we show how REVEAL, a generative AI-guided simulation workflow, can, without any external bias potential, capture the ES2 ↔ GS transitions, estimate the relative populations of metastable states, identify experimentally observable ground states using PWCS, characterize transition paths, and determine relative timescales (**Figure 1**). We also identify limitations of generative AI-based sequence-to-tertiary-structure predictors, such as AlphaFold3^31^, ESMFold2^32^, OpenFold3^33^, Boltz-2^34^, and Protenix-v2^35^, in predicting experimental ensembles. We believe our sampling strategy is straightforward and can be extended to other RNAs to capture functionally important conformational dynamics beyond the single-structure paradigm.

**Figure 1.**
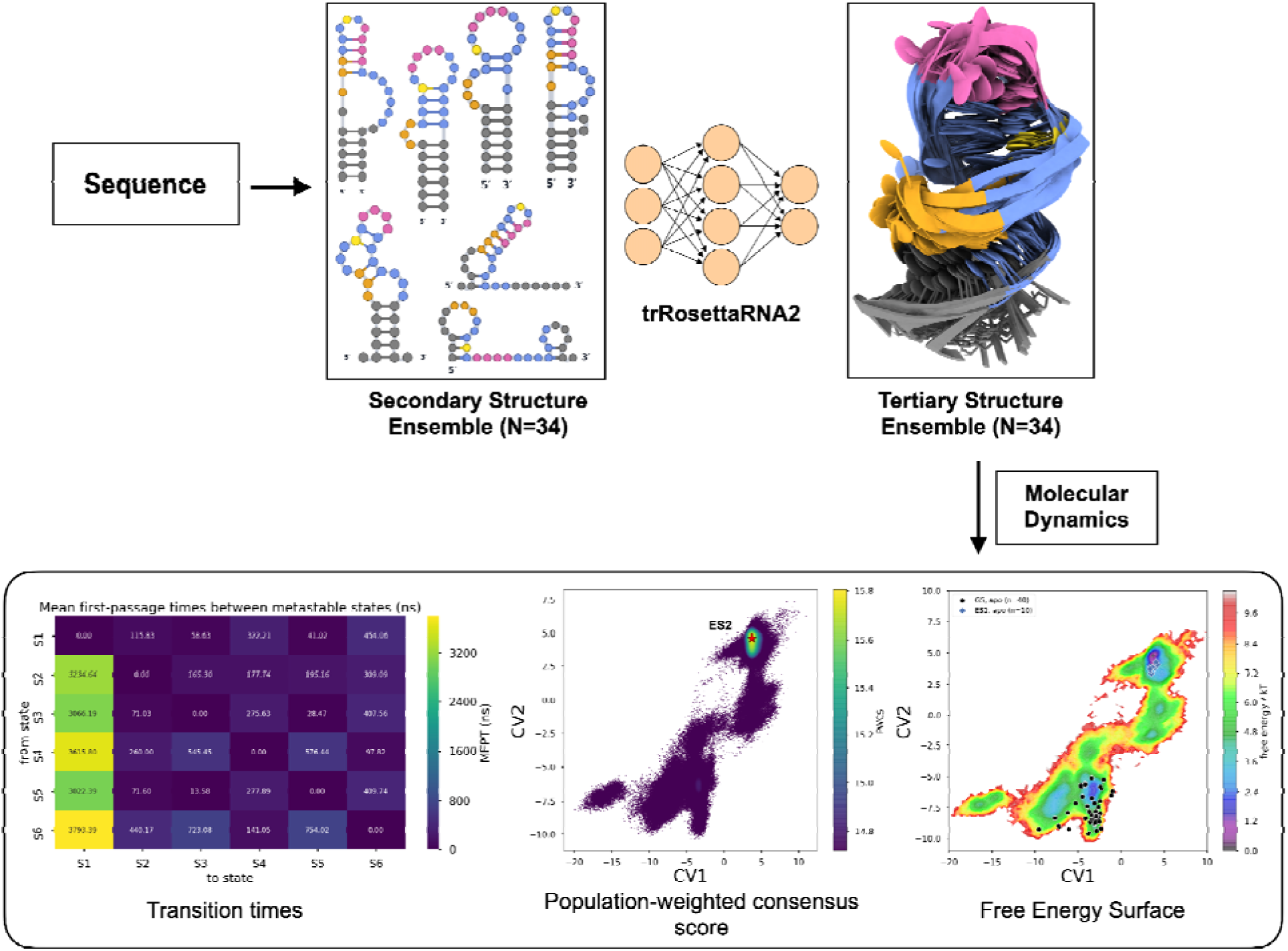
Overview of the RNA Ensemble Validation and Exploration with AI-guided Landscapes (REVEAL) workflow for capturing the thermodynamics and kinetics of RNA conformational heterogeneity in TAR^Mut^. A secondary structure ensemble (N = 34) wa generated from the RNA sequence using ViennaRNA, LinearSampling, and SFold. Representative structures are shown, with nucleotides colored by region: lower stem (gray), UCU bulge (orange), upper helix (blue), UUCG loop (pink), and C13 pivot (yellow). A tertiary structure ensemble (N = 34) was then generated with trRosettaRNA2, conditioned on the secondary structures, and used to initiate multiple independent, unbiased molecular dynamic (MD) simulations. A Markov state model (MSM) connecting these simulations provided Boltzmann weighted ensembles and the kinetics of conformational transitions in TAR^Mut^. The population-weighted consensus score (PWCS), weighted by the MSM, captures the dominant state of TAR^Mut^.

## Results

### REVEAL captures the GS ↔ ES2 transition in HIV-1 TAR^Mut^ beyond NMR-seeded MD and Seq2Ens-MD

TAR^Mut^ was engineered to stabilize ES2 as the dominant state^25^. The ES2 ↔ GS transition is therefore a rare event that conventional MD does not sample without enhanced sampling, making TAR^Mut^ a stringent test for any unbiased physics-based sampling strategy.

We address this with a workflow that combines RNA structural diversity with physics-based molecular simulation. First, ViennaRNA^36^, LinearSampling^37^ and Sfold^38^ predict an ensemble of TAR^Mut^ secondary structures that differ in Watson–Crick pairing. Second, trRosettaRNA2 converts each secondary structure into a tertiary structure, enabling propagation of secondary structure diversity to an ensemble of 34 distinct tertiary structures. Third, each structure in the tertiary structure ensemble seeds an independent unbiased MD simulation (200 ns each; 34 * 200 ns each = 6.8 µs total), and the trajectories are combined using a Markov state model (MSM). Seeding from the structurally diverse initial seeds, rather than from a single experimental structure, enables sampling of the ES2 and GS transitions and captures full spectrum of conformational heterogeneity in TAR^Mut^ without any bias potential. We call this workflow (**Figure 1**) RNA Ensemble Validation and Exploration with AI-guided Landscapes (REVEAL) and describe the GS ↔ ES2 exchange with two collective variables (CVs)^39^:

a. CV1 tracks the pivot base C13, which pairs with G22 in the GS register and with G20 in the ES2 register. CV1 is the C13 N3–G22 N1 distance minus the C13 N3–G20 N1 distance.
b. CV2 tracks the base-pairing register of the helix between the UCU bulge and the UUCG loop. It is the mean Watson–Crick (N1/N3) distance of the four GS-defining pairs (10– 25, 11–24, 12–23, 13–22) minus that of the six ES2-defining pairs (9–24, 10–23, 11–22, 12–21, 13–20, 14–19).

For both CVs, negative values indicate a GS-like register and positive values an ES2-like register.

The MSM-weighted free energy surface (FES) captures reversible transition between the ES2 basin (CV1 ≈ 4 Å, CV2 ≈ 5 Å) to the GS basin (CV1 ≈ −3 Å, CV2 ≈ −6 Å) through a series of intermediate basins (**Figure 2A**). Experimental structures of HIV TAR projected onto CV1 and CV2 fall inside the sampled basins: apo GS NMR models (n = 40; PDB 1ANR^20^, 8THV^40^) occupy the GS basin and ES2 models (n = 10; PDB 8U3M^25^) sit at the ES2 minimum. Unbiased MD of TAR^Mut^ seeded from the ES2 NMR ensemble (NMR-seeded MD) remain stuck in the ES2 basin (**Figure 2B**), despite ∼1.5-fold more aggregate sampling (10 models × 5 replicas × 200 ns; 10 µs).

**Figure 2.**
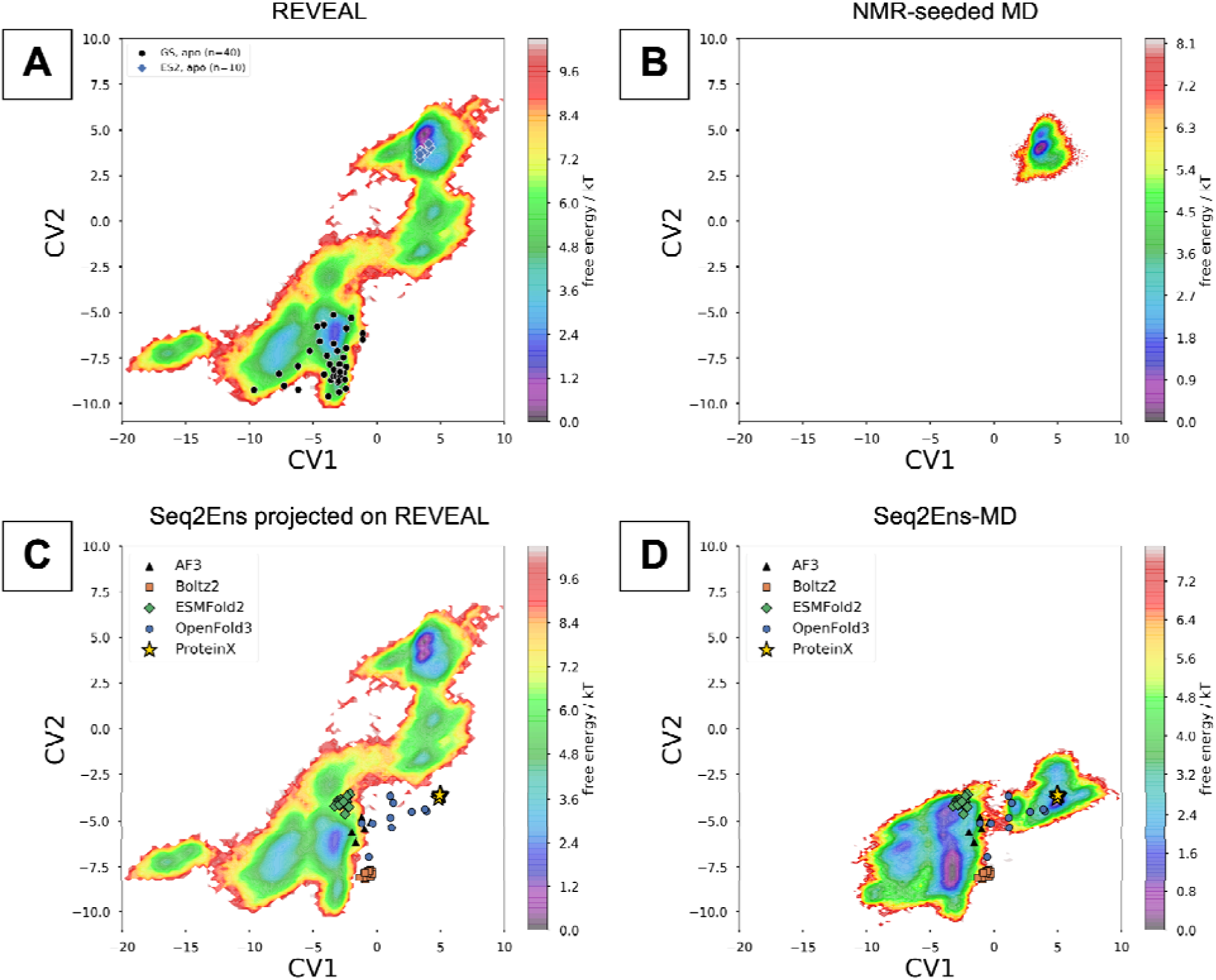
REVEAL samples the full spectrum of conformational heterogeneity in TAR^Mut^, compared to NMR-seeded MD and Seq2Ens-MD. **(A)** MSM-weighted FES projected along CV1 and CV2 (both in Å) from REVEAL (34 replicas × 200 ns; 6.8 µs total) samples the GS, ES2 and intermediate states and identifies ES2 as the dominant basin. Apo NMR models of GS (*black circles*, n = 40; PDB 1ANR, 8THV) and ES2 (*blue diamonds*, n = 10; PDB 8U3M) are superimposed and fall within sampled basins. **(B)** FES from NMR-seeded MD (10 NMR models × 5 replicas × 200 ns; 10 µs total). The simulations remain confined to the ES2 basin and sample neither GS nor the intermediates. **(C)** CV1 and CV2 of structures predicted by the Seq2Ens methods AlphaFold3 (N = 5), Boltz-2 (N = 20), ESMFold2 (N = 20), OpenFold3 (N = 10) and Protenix-v2 (N = 5), projected onto the REVEAL FES from panel A. None of the predicted structure samples the ES2 basin. **(D)** MSM-weighted FES from Seq2Ens-MD started from the structures in panel C. GS appears as the dominant state and ES2 is not sampled, which inverts the experimental population ordering of TAR^Mut^, where ES2 is dominant.

The sampling acceleration achieved by REVEAL is highly dependent on the initial diversity of the tertiary structure ensemble. In REVEAL, conditioning trRosettaRNA2 on the secondary structure ensemble places tertiary structures in the GS basin, the ES2 basin and the intermediate regions between them (**Figure 3A**). Trajectories started from intermediate structures transitions to either GS or ES2 basin within a single 200 ns run: three transitions to ES2, one to a partial ES2 state and two to GS, one of which continues into the GS sub-state S1 (**Figure 3B**). The sampling of GS and ES2 basins from the intermediate states enable generation of a fully connected MSM.

**Figure 3.**
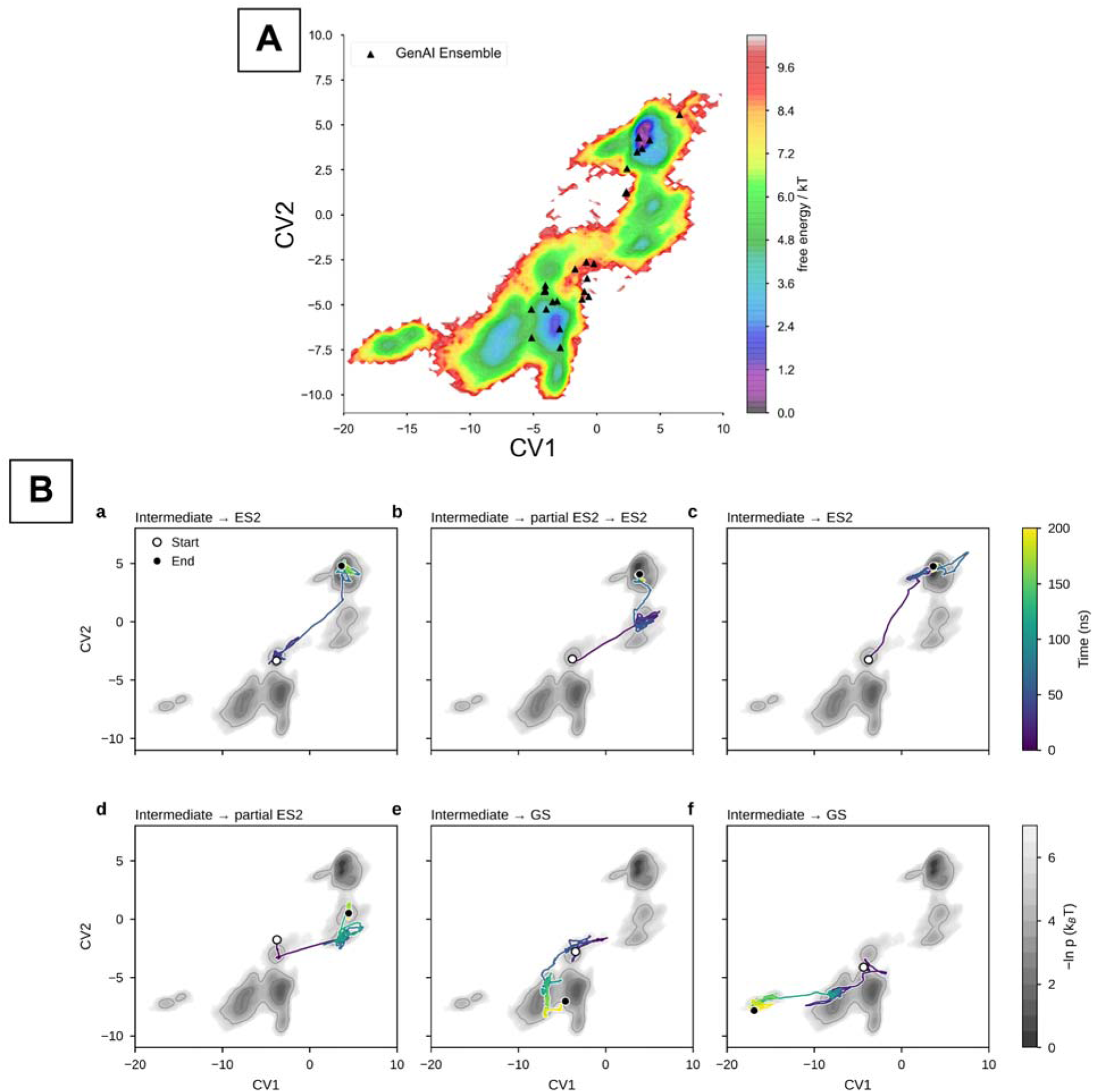
REVEAL seeds sample the GS, ES2 and intermediate regions and yield transitions to both end states. **(A)** trRosettaRNA2 tertiary structures built from the secondary structure ensemble (*black triangles*) projected onto the MSM-weighted free energy surface along CV1 and CV2 (both in Å). The seeds lie in the GS basin, the ES2 basin and the intermediate region between them. **(B)** Representative 200 ns trajectories (a to f) started from intermediate seeds, projected onto CV1 and CV2 and colored by simulation time. Open and filled circles mark the start and end of each trajectory. The gray background shows the MSM-weighted free energy surface (−ln *p*, in *k*_B_*T*). Trajectories a to c reach ES2 (b passes through a partial ES2 state), d reaches a partial ES2 state, and e and f reach GS, with f ending in the S1 region.

Sequence-to-ensemble predictors such as AlphaFold3, Boltz-2 etc. also generate TAR^Mut^ structural ensembles, so we benchmarked them as an alternative seed source. We refer to these models as Seq2Ens predictors and to MD seeded from them as Seq2Ens-MD. Unlike REVEAL, their ensembles are conditioned on sequence alone, and they failed to generate seeds across the GS ↔ ES2 path. Structures from AlphaFold3 (N = 5), Boltz-2 (N = 20), ESMFold2 (N = 20), OpenFold3 (N = 10) and Protenix-v2 (N = 5) all lie at negative CV2, none near the ES2 basin (**Figure 2C**). Boltz-2 places TAR^Mut^ in the GS basin, AlphaFold3 and ESMFold2 near the GS/S2 boundary, and OpenFold3 and Protenix-v2 at positive CV1 but negative CV2, a region the REVEAL FES barely samples. Seq2Ens-MD makes GS the dominant state, forms an off-pathway basin near CV1 ≈ 5 Å, CV2 ≈ −4 Å, and never samples ES2 (**Figure 2D**).

Across NMR-seeded MD, Seq2Ens-MD and REVEAL, the seed ensemble, not the aggregate simulation time, determines whether the GS ↔ ES2 exchange is sampled. NMR-seeded MD remains in the ES2 state from which it starts. Seq2Ens-MD also starts from an ensemble, but its seeds cluster in a single region and carry the predictor’s bias into the dynamics: ES2 is never sampled, and GS is incorrectly predicted as the dominant state. Ensemble size alone is therefore insufficient; what conditions the ensemble matters. RNA register shifts are discrete rearrangements of base pairing, and a secondary structure ensemble samples the relevant registers directly. Conditioning tertiary structure prediction on this ensemble places seeds on both sides of the barrier and in the intermediate region. Unbiased MD connects these seeds into a network of transitions, from which an MSM estimates Boltzmann-weighted populations and transition timescales.

To assess the sensitivity of the MSM to its construction parameters, we varied the number of cluster centers (k = 50, 100 and 200; **Figure 4A**) and the lag time (τ = 10, 100, 200 and 300 frames; **Figure 4B**). In all cases, the MSM-weighted FES preserves the positions of the metastable basins and their relative free energies. The equilibrium description of TAR^Mut^ is therefore insensitive to the choice of state discretization and lag time. Implied timescales also suggest a relative convergence of REVEAL (Figure S1A in Supporting Information).

**Figure 4.**
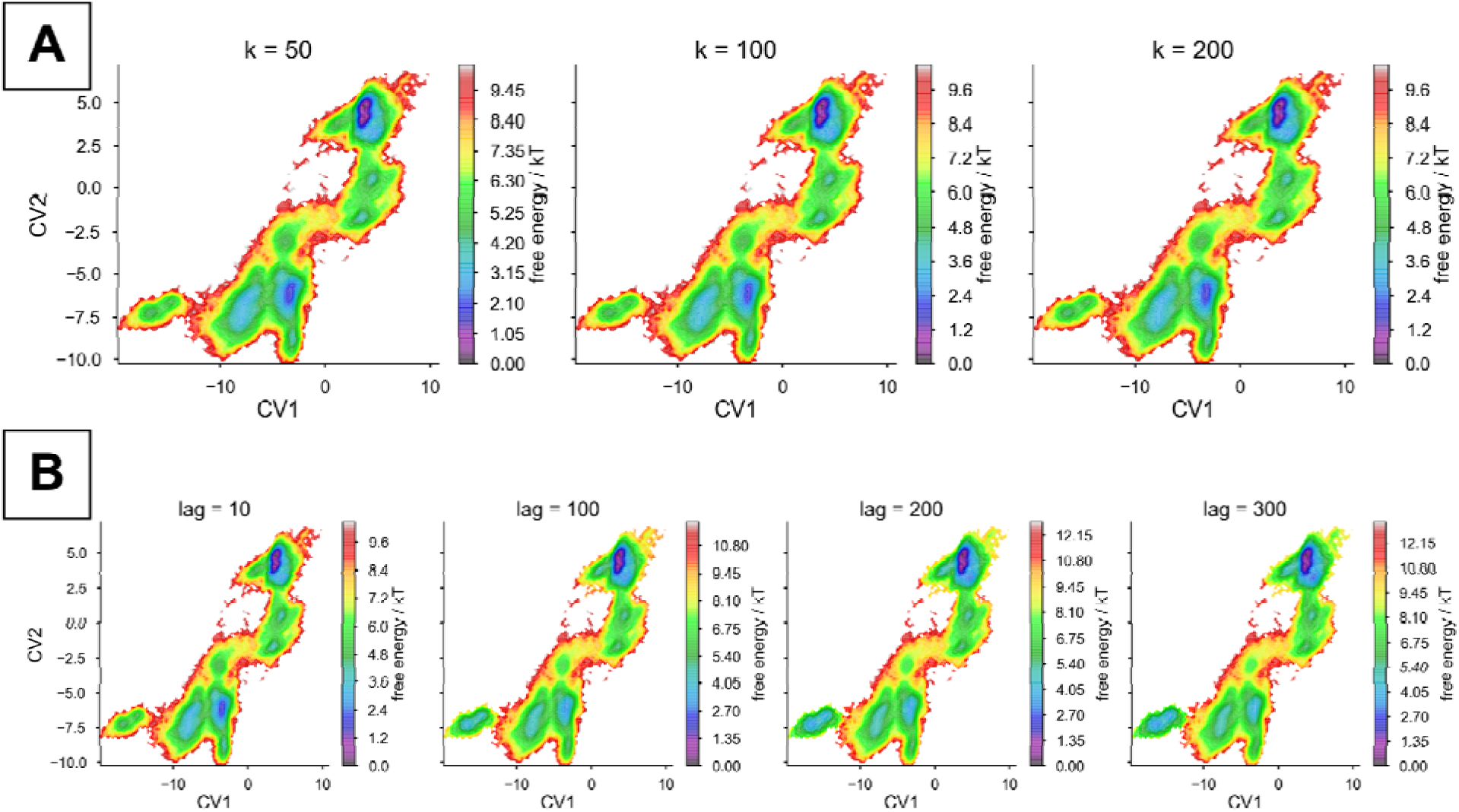
**Convergence of the MSM-weighted FES projected along collective variables CV1 and CV2**. The robustness of the FES is demonstrated across (A) varying numbers of KMeans cluster centers (K=50, 100, 200) and (B) different lag times (10, 100, 200, and 300 frames). The overall shape and relative free-energy values of the metastable basins remain invariant to these parameter choices, indicating apparent convergence of the REVEAL sampling of TAR^Mut^.

### REVEAL resolves thermodynamics and kinetics of the TAR^Mut^ conformational heterogeneity

Macrostate generated using PCCA+ algorithm using CV1 and CV2 feature space yields six metastable states (**Figure 5A**). S6 is ES2 and projection of ES2 NMR models confirm that. The GS basin splits into S3 and S5, which sampled the apo GS NMR models, and S1, a newly identified GS-like sub-state at strongly negative CV1 (≈ −20 to −13 Å). S2 and S4 lie between the two basins.

**Figure 5.**
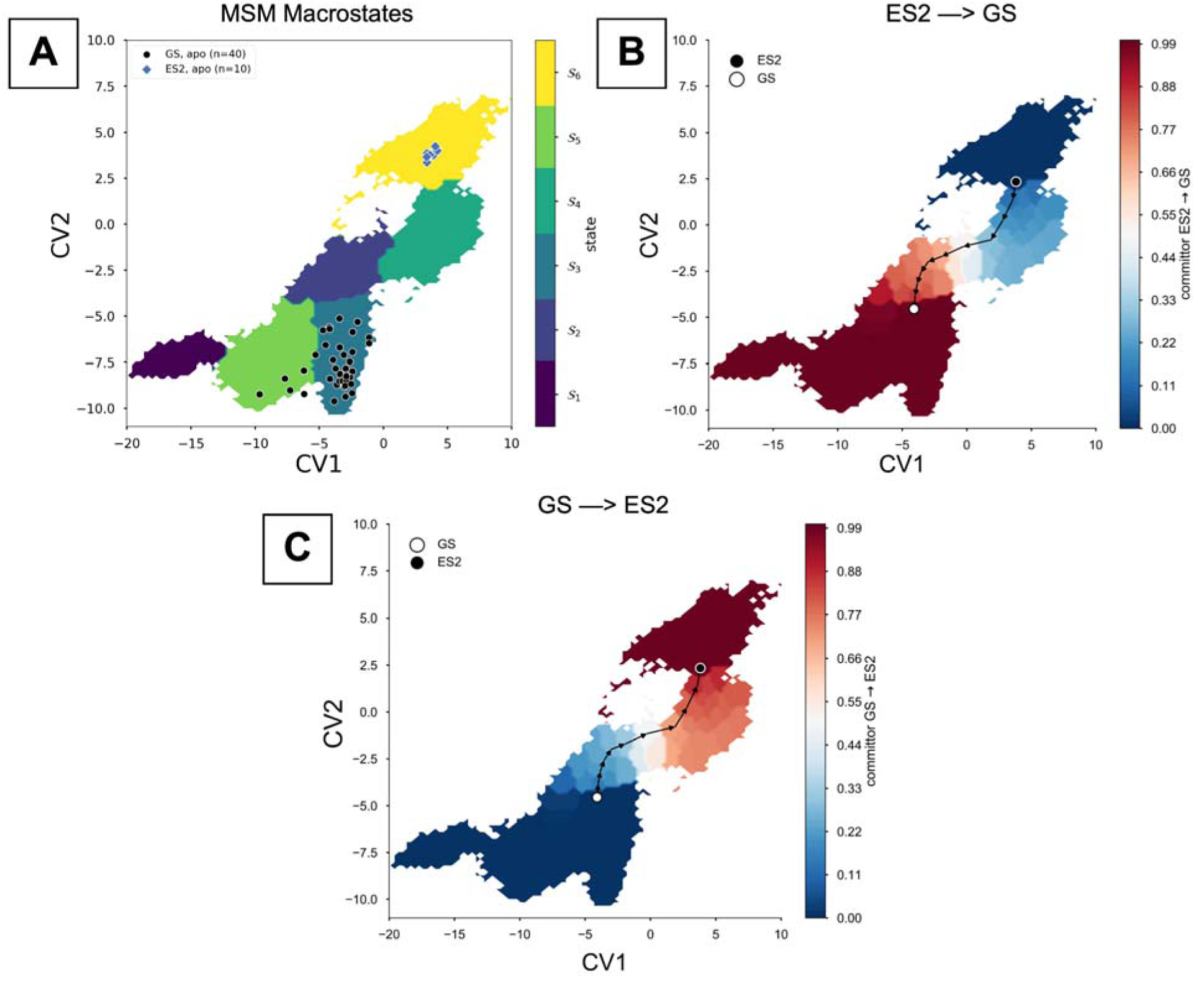
MSM macrostates and committor analysis for the ES2. ↔ **GS transition in TAR^Mut^. (A)** Six metastable states (S1 to S6) from PCCA+^41^ in CV1/CV2 space (both in Å). S6 is ES2 and S3 is GS. Apo NMR models of GS (*black circles*, n = 40; PDB 1ANR, 8THV) and ES2 (*blue diamonds*, n = 10; PDB 8U3M) are superimposed. **(B)** Committor analysis for the ES2 → GS transition, the probability of reaching GS (*open circle*) before returning to ES2 (*filled circle*). The transition path (*black line*) crosses the committor ≈ 0.5 region through the intermediate states (S2 and S4). **(C)** Committor for the reverse transition, GS → ES2, obtained by exchanging the source and sink states. The GS → ES2 and ES2 → GS transition paths pass through the same intermediate states, indicating a single, shared route between the two basins.

The committor analysis identifies S2 and S4 as the intermediates of the ES2 ↔ GS transition (**Figure 5C, D**). The dominant path leaves ES2 through S4, crosses the committor ≈ 0.5 region at the S4/S2 boundary and transitions to GS through S2. MFPTs between metastable states are consistent with this landscape. The S4 → S6 transition occurs within 98 ns, whereas transitions from S4 to the GS sub-states (S4 → S3, S4 → S5) proceed on a slower timescale of 546–576 ns, placing S4 on the ES2 side of the transition barrier. Conversely, transitions from S2 to the GS sub-states (S2 → S3, S2 → S5; 165–195 ns) are faster than the S2 → ES2 transition (309 ns), placing S2 on the GS side. The GS → ES2 transition is faster than the reverse (S3 → S6: 408 ns; S6 → S3: 723 ns). Within the GS basin, S3 and S5 interconvert on a 14–28 ns timescale. Transitions into S1 are slow, with MFPTs of 3.0–3.8 µs from all other states, whereas transitions out of S1 to S5 and S3 occur within 41 and 59 ns, respectively, identifying S1 as a transiently populated GS sub-state (Figure 7; 95% confidence intervals of the MFPTs are given in Figure S2 in Supporting Information).

The MSM equilibrium populations rank the states S6 (π = 0.589) > S3 (0.209) > S5 (0.128) > S4 (0.045) > S2 (0.015) > S1 (0.014) (Figure 6; 95% confidence intervals in Figure S1B in Supporting Information). ES2 (S6) is about 1.7-fold more populated than the entire GS basin (S1 + S3 + S5, π = 0.351), a free energy difference of ∼0.5 kT, consistent with the faster GS → ES2 than ES2 → GS kinetics and the experimental observation that ES2 is the dominant state in TAR^Mut^.

**Figure 6.**
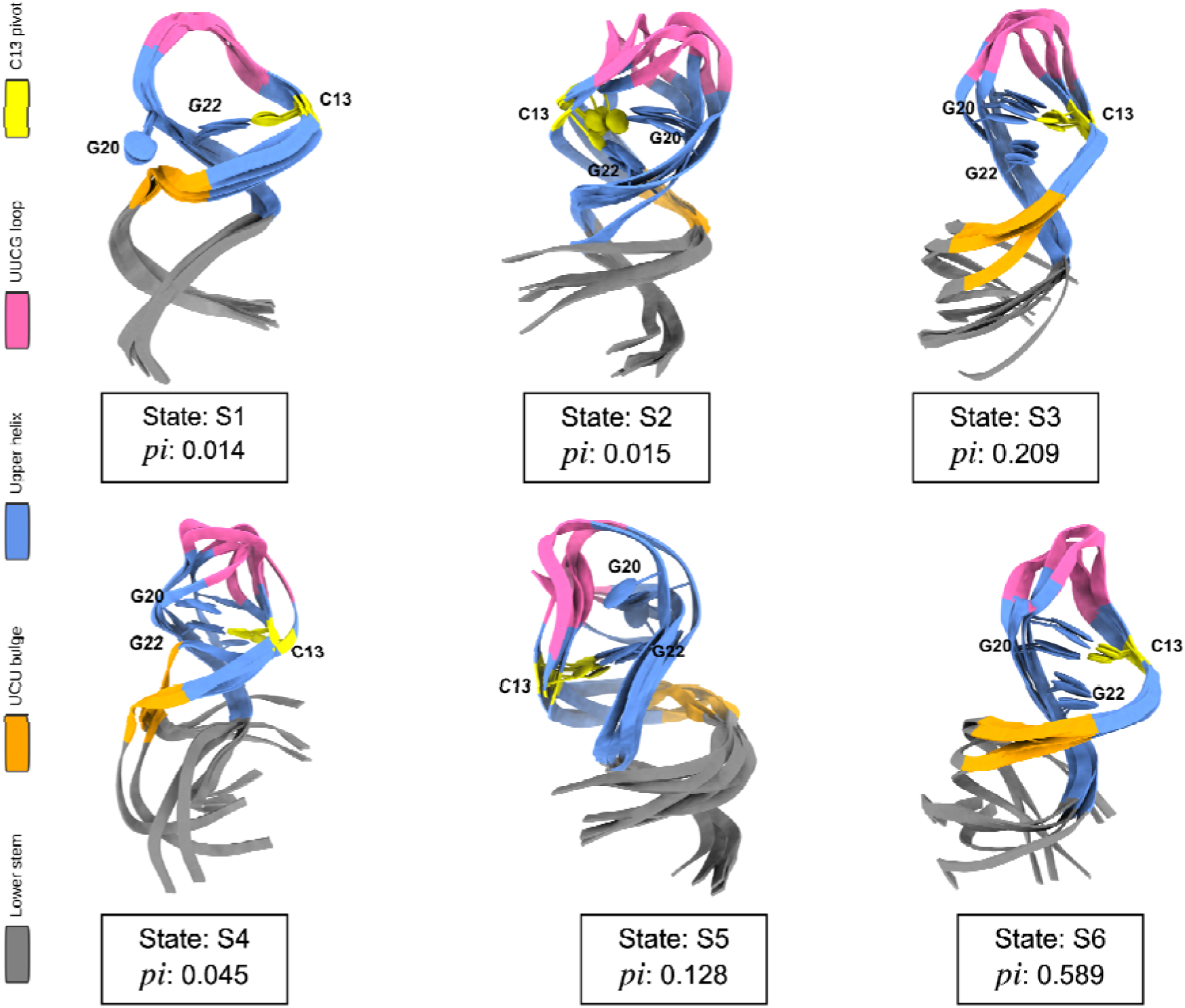
Representative structures of the six metastable states of TAR^Mut^. For each state (S1 to S6) identified by PCCA+ in CV1/CV2 space, the five structures with the highest PWCS among 20 representative structures are overlaid. The upper helix (*blue*) is closed by the UUCG apical loop (*pink*) and connected to the lower stem (*gray*) through the UCU bulge (*orange*). C13 (*yellow*) is shown relative to G20 and G22 to illustrate its pivot between the GS (G22) and ES2 (G20) pairing registers, together with the conformational heterogeneity of the lower stem, UCU bulge, upper helix and apical loop. MSM equilibrium populations (π) are given for each state. ES2 (S6, π = 0.589) is the dominant state of TAR^Mut^ and is about 1.7-fold more populated than the entire GS basin (S1 + S3 + S5, π = 0.351).

**Figure 7.**
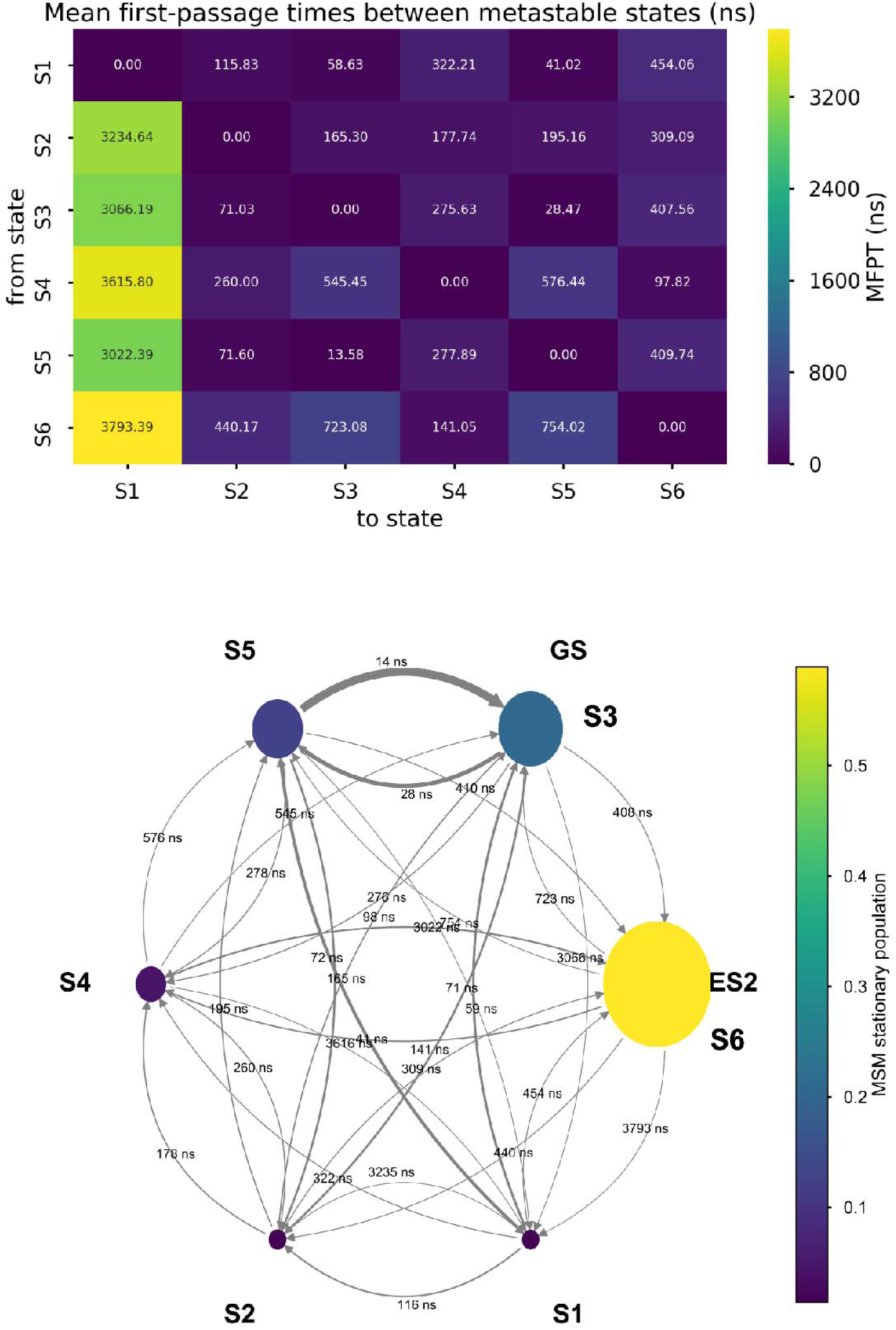
Kinetics of the metastable states of TAR^Mut^ from the MSM. Upper panel: Mean first-passage times (MFPTs, in ns) between the six metastable states (S1 to S6). Rows give the initial state and columns the final state. **Lower panel:** Network representation of the MSM. Node area and color give the MSM stationary population of each state. ES2 (S6) is the most populated state, followed by the GS sub-states S3 and S5, and S1 is a newly identified GS sub-state. Arrows give the direction of each transition, are labeled with the MFPT in *ns*, and are drawn thicker for shorter MFPTs. The GS → ES2 transition is faster than the reverse (S3 → S6: 408 ns; S6 → S3: 723 ns). S3 and S5 interconvert within 14 to 28 ns, whereas transitions into S1 are slow (3.0 to 3.8 µs from any other state).

Additionally, we scored each frame with a population-weighted consensus score (PWCS): the mean normalized log-likelihood from Gaussian mixture models (2–8 components) fit to CV1 and CV2. The models are fit to a resample weighted by the MSM equilibrium populations, so frame are scored against the Boltzmann distribution rather than the raw sampling. PWCS peaks sharply in the ES2 register, with only a faint secondary maximum in the S3 basin (**Figure 8**) and independently recovers ES2 as the dominant state. The five top-ranked PWCS structures of each state show the C13 pivot between G20 and G22 along with heterogeneity in the lower stem, UCU bulge, upper helix and UUCG loop (**Figure 6**). These state-resolved, population-weighted structures are directly usable for downstream state-specific virtual screening or binder design.

**Figure 8.**
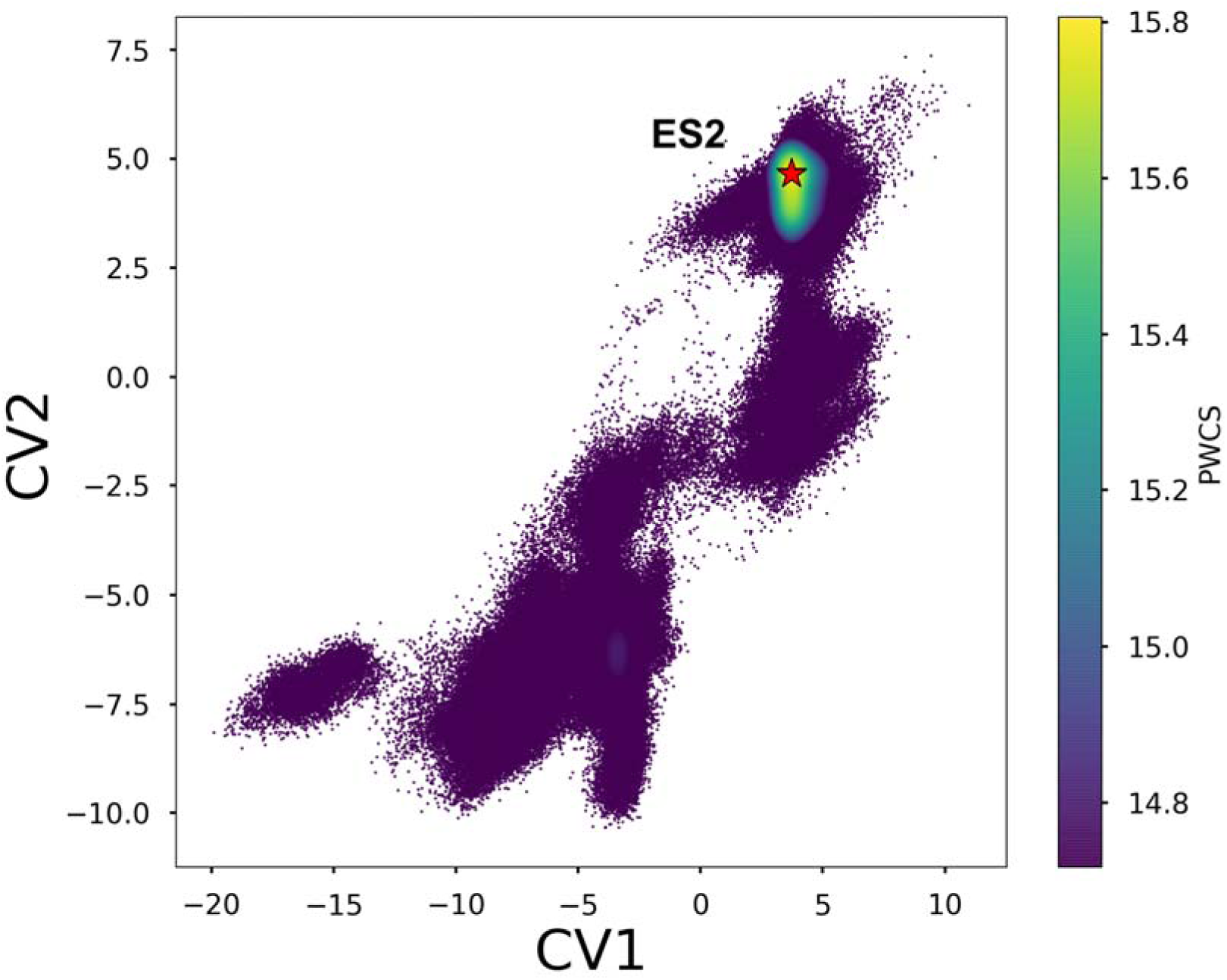
Population-weighted consensus score (PWCS) identifies ES2 as the dominant state of TAR^Mut^. PWCS was calculated for every frame and plotted on the CV1/CV2 plane (both in Å), with each point colored by its MSM-weighted PWCS. PWCS is the mean normalized log-likelihood from Gaussian mixture models (2 to 8 components) fit to CV1 and CV2 on a resample weighted by the MSM equilibrium populations, so higher values mark conformers that are more representative of the Boltzmann-weighted ensemble. The color scale is clipped at the median to emphasize the high-scoring region. PWCS peaks in the ES2 basin (*red star*), with only a weak secondary maximum in the GS basin.

### Functional PCA outperforms time-lagged learned CVs in resolving TAR^Mut^ states

Conceptually simple CV1 and CV2 to describe conformational heterogeneity of TAR^Mut^ also serves as a benchmark for data-driven dimensionality reduction methods. We applied four methods to the G-vector representation of TAR^Mut^: functional principal component analysis (FPCA), slow feature analysis (SFA)^17^, time-lagged autoencoder (TAE)^42,43^ and deep time-lagged independent component analysis (DeepTICA)^44^. Only the first FPCA component resolves all three regions, positive in GS, negative in ES2 and near zero in the intermediates (**Figure 9A**). The TAE latent separates GS from the rest but assigns intermediates the same sign as ES2 (**Figure 9C**). SFA and DeepTICA remain near zero over most of the surface, with contrast confined to scattered ES2 bins (**Figure 9B, D**). This highlights that no dimensionality reduction method is reliable by default for capturing conformational heterogeneity of RNA register shifts. Learned CVs should be checked against a physically interpretable reference before they are used for MSM featurization or as biasing coordinates.

**Figure 9.**
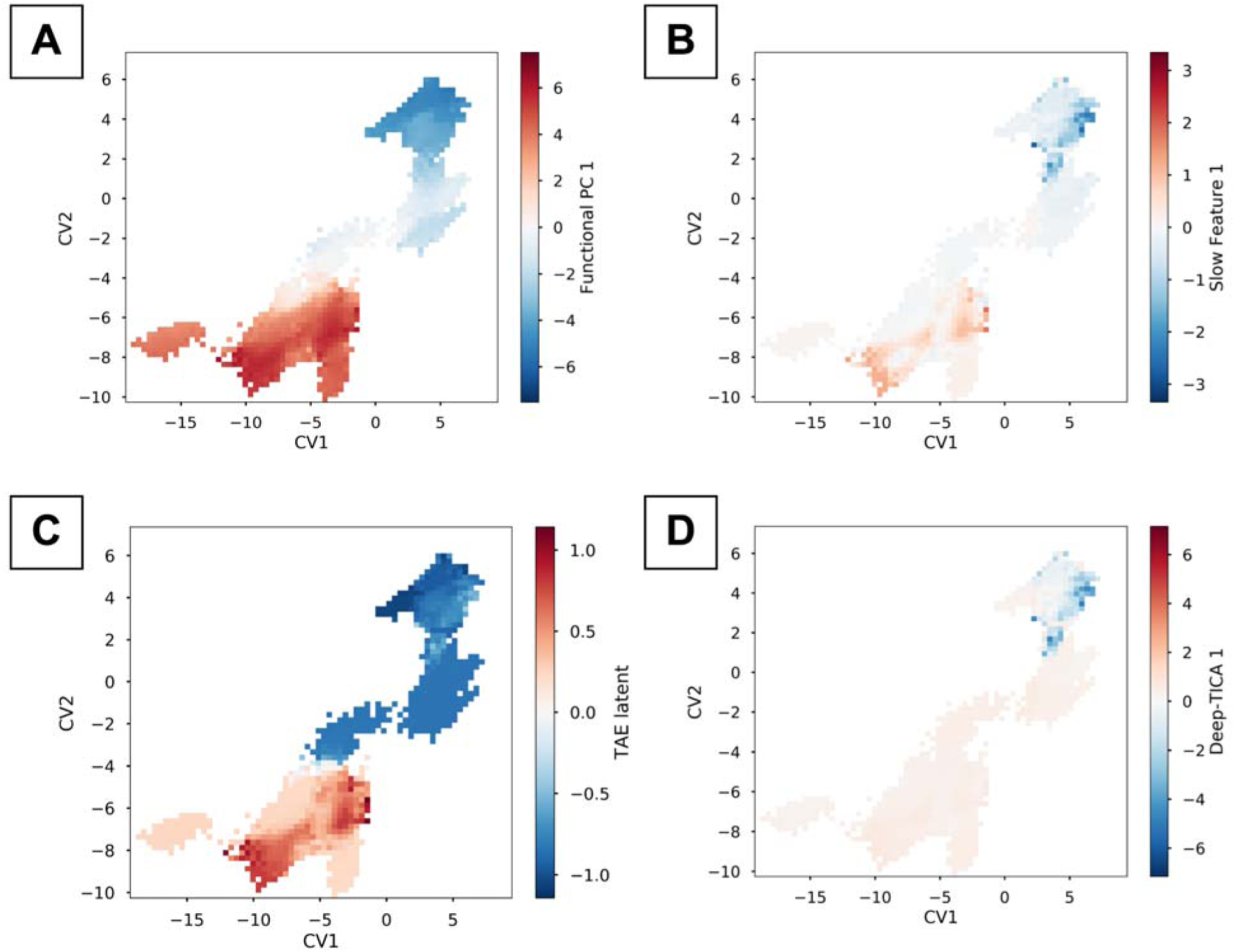
Comparison of dimensionality reduction methods for the G-vector representation of TAR^Mut^. The first component of each method is shown on the CV1/CV2 plane (both in Å), with each bin colored by the component value. **(A)** Functional PCA separates GS (positive values), ES2 (negative values) and the intermediate states (values near zero). **(B)**Slow feature analysis (lag = 10), **(C)** time-lagged autoencoder (TAE; lag = 10) and **(D)** DeepTICA (lag = 10) do not resolve all three regions. SFA and DeepTICA are near zero over most of the FES, whereas TAE separates GS from ES2 but assigns the intermediates the same sign as ES2. These results indicate that the choice of dimensionality reduction method affects how well the metastable states of an RNA are resolved.

As an independent check, we rebuilt the MSM in the latent spaces of FPCA, SFA, TAE and DeepTICA and projected the resulting MSM-weighted populations onto the CV1/CV2 plane. The relative populations of the metastable states are preserved across all four representations, indicating that the state definitions and the population ordering of TAR^Mut^ do not depend on the choice of featurization. Importantly, PWCS itself does not depend on the choice of collective variables: when recomputed in the FPCA, SFA, TAE and DeepTICA latent spaces, PWCS recover ES2 as the dominant state (Figures S3 and S4, Supporting Information), confirming that the population-weighted ranking is robust to the underlying featurization.

## Conclusions

In this work, we introduced an intuitive workflow that combines secondary structure ensemble generation, secondary structure–conditioned tertiary structure ensemble prediction with trRosettaRNA2, unbiased MD and Markov state modeling to sample the reversible ES2 ↔ GS register shift in TAR^Mut^, a rare conformational transition beyond the reach of traditional unbiased molecular dynamics simulation. With 6.8 µs of aggregate sampling distributed across 34 seeds, REVEAL recovers the experimentally observed GS and ES2 basins, resolves two on-pathway intermediates (S2 and S4) and a new GS sub-state (S1), and identifies ES2 as the dominant state of TARMut, consistent with the design objective of the construct. By contrast, ∼10 µs of MD seeded from the ES2 NMR ensemble remains trapped in the ES2 basin, and MD seeded from sequence-only ensemble predictors (AlphaFold3, Boltz-2, ESMFold2, OpenFold3 and Protenix-v2) never samples it. To our knowledge, this is the first Markov state model of TAR^Mut^, and it provides a state-specific description of the populations, transition pathways and relative timescales of the ES2 ↔ GS exchange, which has so far been accessible only through enhanced sampling with external bias potentials.

The central lesson is that, for RNA register shifts, whether a transition is sampled in REVEAL depends on the tertiary structure ensemble used to seed the simulations, not on the aggregate simulation time. Register shifts are discrete rearrangements of base pairing, and hence the relevant degrees of freedom can be enumerated in secondary structure space, where thermodynamic models are inexpensive and comparatively well calibrated. Conditioning tertiary structure prediction on this ensemble turns an intractable search over tertiary conformations into a tractable one: the generative model places structures on both sides of the barrier and along the intermediate states, physics-based sampling connects them, and the MSM provides Boltzmann-weighted state populations and transition times. This logic is equivalent to recent strategies for globular proteins, in which structural ensembles generated by MSA subsampling, diffusion or flow-matching models seed unbiased simulations that are subsequently integrated through MSMs or related frameworks, as in AlphaFold2-RAVE^13^, AlphaFold-MSM^15^, BioEmu-MSM^16^ and AlphaFold-SFA^17^. REVEAL extends this paradigm to RNA, where the natural source of structural diversity is the secondary structure ensemble rather than a coevolutionary or learned sequence–structure prior. In both settings, the generative model is not asked to predict the ensemble, only to generate structures that serve as seeds; populations and kinetics come from the choice of force field and water model and from the statistical protocols used to analyze the aggregate simulation data. We regard this separation as the main reason such approaches are robust, and more defensible use of generative AI in molecular simulation rather than treating predicted structural ensembles as thermodynamic ensembles in their own right.

The behavior of the sequence-only ensemble predictors reinforces this point. All five predictors place every TAR^Mut^ structure in intermediates that lie off the ES2 ↔ GS transition pathway and close to the GS-like register, even though the construct was engineered to favor ES2. MD launched from these structures inherits this bias and inverts the experimental population ordering. A plausible interpretation is that these models, trained on structural data in which RNA is sparsely represented and dominated by single ground-state conformers, default to conformations near the most-represented fold of the wild-type sequence family and respond only weakly to the few mutations that shift populations. These observations caution against using co-folding models as drop-in seed generators for RNA dynamics without first verifying that their outputs span the relevant states.

A further implication concerns how RNA ensembles are determined when experimental data are limited. Ensemble descriptions of HIV-1 TAR have so far relied on NMR data, most prominently residual dipolar couplings, to select or reweight conformers from a pre-generated FARFAR2 library^25^. This strategy is powerful when such data exist, but it presupposes them, and for TAR^Mut^, as for most RNAs of therapeutic interest, comparable NMR datasets are unavailable or cover only a subset of states. REVEAL combined with PWCS offers a route that does not depend on prior experimental restraints. The MSM supplies Boltzmann weights from physics-based dynamics, and PWCS uses those weights to rank individual conformers by how representative they are of the equilibrium ensemble; in TAR^Mut^, this recovers ES2 as the dominant state, with the GS register as a secondary maximum, without any reweighting against NMR residual dipolar coupling data. Crucially, PWCS does not require a low-dimensional representation that separates the metastable states crisply: because it scores conformers directly against the MSM-weighted Boltzmann distribution, it identifies the dominant state even when learned collective variables fail to resolve the states (Figures S3 and S4, Supporting Information). This is what distinguishes the approach from prior ensemble descriptions, which identify the relevant states only indirectly, by back-calculating NMR residual dipolar couplings from a pre-generated library and matching them to experiment; PWCS removes that dependence. We do not view this as a replacement for experiment. Rather, MSM-weighted PWCS provides testable structural hypotheses and a physically weighted prior ensemble that NMR data, when they become available, can refine, and it extends ensemble-based structural analysis to RNAs for which such data do not yet exist.

The statistical model underlying these results for TAR^Mut^ is robust: the REVEAL simulations are converged, the MSM-weighted free energy surface is insensitive to the number of cluster centers and the lag time, and Bayesian MSM estimation enables prediction of statistical uncertainties corresponding population and timescale estimates. The principal caveats therefore concern the physical model rather than the statistical one. The populations and transition times reported here are properties of the chosen force field, water model and ionic conditions as much as of TAR^Mut^ itself. Together these choices can shift the relative population and timescales of conformational states and the barriers between them. Ionic conditions add a further layer. Counterions screen the highly charged phosphate backbone, and divalent ions such as Mg^2+^ can bind site-specifically at bulges and loops, so ion type, concentration and ion parameters can redistribute register populations and alter the barriers that separate them^45,46^. How force fields^47,48^, water models^49–51^, counterions and ion types shape the populations and timescales recovered by REVEAL, remains an active area of research, and the ∼0.5 kT ES2–GS free energy difference should therefore be read as qualitative agreement with the design of TAR^Mut^, in which ES2 is the dominant state, rather than as a quantitative prediction. The kinetics warrant even greater caution. To our knowledge, the ES2 ↔ GS exchange rate has not been measured for the 31-nucleotide TAR^Mut^ construct; the millisecond lifetime of ES2 was determined by relaxation dispersion NMR on the 29-nucleotide wild-type TAR, in which ES2 is only sparsely populated. Because the mutations that make ES2 dominant in TAR^Mut^ also reshape the relative stabilities of the two registers, and very likely the barrier between them, the wild-type rates cannot serve as a direct benchmark, and the MSM timescales reported here should be regarded as predictions rather than validated quantities. Water models such as TIP3P that underestimate solvent viscosity accelerate diffusive dynamics, and ion parameters that misrepresent ion–phosphate binding affinities or ion exchange rates can further alter the effective barriers to base-pair rearrangement. We therefore expect the ordering of timescales, such as the faster GS → ES2 than ES2 → GS transition and the transient character of S1, to be more robust than their absolute magnitudes. REVEAL addresses the sampling problem, and its outputs will improve as force fields, water models and ion parameters improve.

Two further caveats regarding the choice of CVs and the scoring scheme. First, CV1 and CV2 encode prior knowledge of the GS and ES2 registers. This is appropriate for a benchmark system, but for RNAs whose alternative registers are unknown, the featurization must be learned, and our comparison shows that time-lagged learned CVs (SFA, TAE and DeepTICA) do not reliably represent the GS, ES2 and intermediate states of TAR^Mut^, whereas FPCA does. This behavior is case dependent, so for every system it is important to check which low-dimensional representation does a better job of discriminating states before those CVs are used to perform MSM or biasing coordinates for enhanced sampling. Second, PWCS does not depend on the choice of CVs. We have shown that PWCS can correctly identify the dominant state in TAR^Mut^ even when the low-dimensional representation fails to crisply identify that state. This makes PWCS a generalizable method for identifying dominant states in diffusive systems such as RNA and intrinsically disordered proteins, where constructing a correct low-dimensional representation for the necessary downstream post-processing is often difficult. Further, MSM reweighting is not intrinsic to PWCS. The same consensus scoring can be applied directly to energies obtained from simulations, which would extend it to systems for which a fully connected MSM is not available. We are exploring this energy-based formulation of PWCS in upcoming studies.

More broadly, this work argues for a measured view of what generative AI contributes to RNA conformational sampling. The gains reported here come not from a new algorithm but from a transparent combination of open-source tools, each with a clearly defined role. We suggest that descriptions of such pipelines state clearly where structural information enters, whether from secondary structure models, tertiary structure predictors or the force field, because this determines both what a method can sample and how its failures should be diagnosed.

Natural extensions include systematic benchmarking of REVEAL across RNA force field, water model and ionic condition combinations, adaptive reseeding from under-sampled MSM states, and quantitative validation against NMR observables such as residual dipolar couplings and chemical shifts, with ensemble reweighting where appropriate. Application of REVEAL to riboswitches^52,53^ and other RNAs whose functional states involve tertiary contacts, pseudoknots^54^ or Mg^2+^-dependent folding^55^ will test the limits of seeding from a secondary structure ensemble, which may not fully capture such features. Because REVEAL decouples seed generation from the physical model, it also offers an efficient testbed for force field development: the same seed ensemble can be propagated under different parameter sets to compare their predicted populations and timescales against experiment. For HIV-1 TAR, the state-resolved, population-weighted structures of ES2 and the intermediates provide a starting point for conformation-selective screening against the auto-inhibited state. More generally, moving beyond the single-structure paradigm in RNA will require coupling inexpensive generators of structural diversity with physical models that can assign that diversity its correct thermodynamic and kinetic weight^4^.

### Computational Methods Ensemble Generation

The starting pool of conformations was generated from an ensemble of secondary structures predicted for the TAR^Mut^ sequence. Non-redundant Boltzmann sampling with ViennaRNA^36^ was used to draw secondary structures from the equilibrium distribution, which were clustered by base-pair distance into 20 representative structures. A further 10 and 4 secondary structures were generated with LinearSampling^37^ and Sfold^38^, respectively. Each of these secondary structures was converted into a three-dimensional model with trRosettaRNA2^5^, conditioned on the corresponding secondary structure, and one additional model was predicted with trRosettaRNA2 from sequence alone, giving 35 candidate tertiary structures. Each candidate was then screened to confirm that the base pairs formed in the three-dimensional model matched those of its target secondary structure; one structure that failed this check was discarded, leaving 34 seeds for molecular dynamics.

Tertiary structure ensembles were also generated directly from the TAR^Mut^ sequence using Seq2Ens predictors. Structures were predicted with AlphaFold3 (N = 5), Boltz-2 (N = 20), ESMFold2 (N = 20), OpenFold3 (N = 10) and Protenix-v2 (N = 5), giving 60 predicted tertiary structures. Unlike the trRosettaRNA2 models above, these ensembles are conditioned on sequence alone rather than on a predicted secondary structure ensemble. The resulting structures were used as seeds for Seq2Ens-MD.

### Molecular Dynamics Simulations

Each tertiary structure predicted using trRosettaRNA2^5^ was prepared for simulation with *tleap* (AmberTools). The RNA was described by the AMBER OL3 force field (leaprc.RNA.OL3)^56^ and solvated using TIP3P^57^ water model. Each system was placed in a truncated octahedral box with a 10 Å solvent buffer, and Na+ counterions were added to neutralize the net charge with no additional salt. The resulting Amber topology and coordinate files were converted to GROMACS format with an in-house script (*amber2gromacs.py*), and all subsequent simulations were performed in GROMACS^58^.

Following energy minimization, each system was equilibrated for 100 ps in the NVT ensemble at 300 K with harmonic position restraints on the RNA. The restraints were then removed, and each system was equilibrated for a further 200 ps in the NPT ensemble at 300 K, with pressure maintained at 1 bar by the Parrinello–Rahman barostat^59^ and temperature controlled by the v-rescale thermostat.

Production simulations were conducted in the NPT ensemble at 300 K and 1 bar using the leap-frog integrator with a 2 fs timestep. A 1.0 nm cutoff was used for nonbonded interactions, while long-range electrostatic interactions were treated via the Particle Mesh Ewald (PME) method^60^ with a 0.16 nm grid spacing, and covalent bonds to hydrogen atoms were constrained with the LINCS algorithm^61^. Each of the 34 seed structures was simulated independently for 200 ns, yielding 6.8 µs of aggregate sampling, with coordinates saved every 10 ps. All equilibration and production runs were carried out using GROMACS.

The NMR ensemble and the Seq2Ens predicted structures were processed with the identical protocol described above, using the same OL3 force field, TIP3P water model, Na+ neutralization, box setup, energy minimization, NVT and NPT equilibration, and production settings. The three sets differ only in their seed structures and therefore in their total aggregate simulation length. NMR-seeded MD was seeded from 10 ES2 apo NMR models, each simulated in 5 replicas for 200 ns, yielding 10 µs of aggregate sampling. Seq2Ens-MD was seeded from 60 predicted structures (AlphaFold3, N = 5; Boltz-2, N = 20; ESMFold2, N = 20; OpenFold3, N = 10; Protenix-v2, N = 5), each simulated for 200 ns, yielding 12 µs of aggregate sampling, compared with the 6.8 µs of REVEAL (34 seeds x 200 ns).

All residues are reported in the sequential GROMACS numbering (1–31). Residues 1–14, 15–18 and 19–31 correspond to G17–C30, U51–G54 and G33–C45, respectively, in the deposited PDB: 8U3M structure. For example, C13, G20 and G22 correspond to C29, G34 and G36 in PDB: 8U3M.

### Featurization

Two collective variables (CVs) were used to describe the GS ↔ ES2 register shift of TAR^Mut^, both computed with MDTraj^62^ from the Watson–Crick heavy-atom donor and acceptor positions (N1 for the purines G and A, N3 for the pyrimidines C and U). Residues are reported in construct numbering (1–31) for the 31-nucleotide sequence 5′-GGCAGAUCUGAGCCUUCGGGAGCUCUCUGCC-3′.

Each CV was evaluated for every frame of every trajectory, and all distances were converted from nanometers to angstrom. CV1 is a pivot-based coordinate centered on C13, the base that pairs with G22 in the GS register and with G20 in the ES2 register. It is defined as the C13(N3)– G22(N1) distance minus the C13(N3)–G20(N1) distance, so that the partner switch of the single most diagnostic base is captured by one scalar. Negative values report retention of the GS partner and positive values report the ES2 partner.

CV2 is a coarser register coordinate that averages over the full set of base pairs that reorganize between the two registers. It is defined as the mean Watson–Crick (N1/N3) distance of the four GS-defining pairs (10–25, 11–24, 12–23 and 13–22) minus the mean of the six ES2-defining pairs (9–24, 10–23, 11–22, 12–21, 13–20 and 14–19).

For both coordinates, negative values correspond to a GS-like register and positive values to an ES2-like register. CV1 and CV2 are strongly correlated on the experimental validation set (Pearson r > 0.9), confirming that they capture the same register transition at different levels of coarse-graining. The two coordinates were combined into a per-frame (CV1, CV2) feature vector for each trajectory, which served as the input space for the Markov state model described below. The same two CVs were also computed for the trRosettaRNA2 tertiary structure ensemble used to seed REVEAL and for the Seq2Ens predicted structures (AlphaFold3, Boltz-2, ESMFold2, OpenFold3 and Protenix-v2), so that seeds, predicted structures and simulation frames could all be projected onto the same CV1/CV2 surface (**Figure 2**).

To relate the CVs to experimentally observed states, decision bands were calibrated against apo NMR models only; ligand-bound structures were excluded. The apo set comprised 40 GS models (PDB 1ANR, 8THV) and 10 ES2 models (PDB 8U3M). GS models gave CV1 = −4.83 ± 2.62 Å and ES2 models CV1 = +3.70 ± 0.33 Å, from which we defined CV1 ≤ −0.28 Å as the GS register and CV1 ≥ +0.75 Å as the ES2 register.

Beyond the two register coordinates, we computed a higher-dimensional, geometry-aware featurization of base-pair arrangement based on R-vectors and G-vectors, following the G-vector construction of Bottaro, Di Palma and Bussi^63^. Each nucleotide was represented by the center of its six-membered aromatic ring, defined as the mean position of the N1, C2, N3, C4, C5 and C6 atoms. These atom names are shared by purines and pyrimidines, so a single definition applies to every residue without base-type-specific atom selections.

For each pair of nucleotides, (*i,j*)the R-vector is the displacement between ring centers:

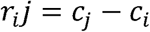

The corresponding G-vector maps this displacement into a smooth four-component representation:

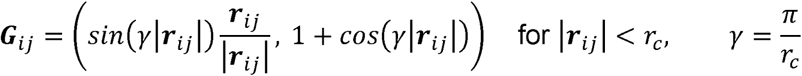

and zero otherwise, so that both the direction and the magnitude of each interbase vector are encoded in a continuous form that tapers to zero at the cutoff. Because *r_ji_ r_ij_* and the first three G-vector components change sign under exchange of *i* and *j* while the fourth (distance-only) component does not, only the *i < j* upper triangle of pairs was retained. All ring centers were computed per frame with MDTraj and distances converted from nanometers to angstrom.

The cutoff *r_c_* was set empirically to 15 Å for this system rather than taken from a literature value, chosen to retain local stacking contacts. Because the *sin* and 1+ *cos* terms taper smoothly to zero at, the hard cutoff discards only the already-negligible tail beyond it. The R and G-vector features were indexed by trajectory and frame in the same order as CV1 and CV2, allowing the feature sets to be combined frame-for-frame in the population-weighted consensus score.

## Markov State Model

Markov state models (MSMs)^18,64–66^ were built with PyEMMA^67^ from the CV1/CV2 featurized trajectories. The conformational space was discretized into 200 microstates by k-means clustering (k = 200), and every trajectory frame was assigned to this microstate representation. Transition matrices were then estimated at a lag time chosen from the convergence of the implied timescales (10 frames, 100 ps). Both maximum-likelihood and Bayesian MSMs were estimated; the Bayesian model additionally provided posterior uncertainties on the stationary and kinetic properties.

To confirm that the results were not artifacts of the discretization, we tested the effect of the number of cluster centers and of the lag time on the MSM-weighted free energy surface and found it to be robust to both choices. The stationary distribution of the MSM provided the equilibrium weights used to construct the free energy surfaces along CV1 and CV2 (computed as *−kT ln P*) and to population-weight the consensus score. Metastable states were identified by coarse graining the microstates with PCCA+^41^, and the interconversion kinetics between them were characterized by mean first passage times.

## Population weighted consensus scoring

To identify conformers that are representative of the equilibrium ensemble, we introduced “*population-weighted consensus score* (PWCS)”. PWCS assigns each frame a single score that reflects how well that conformer is supported across multiple feature representations and density models of varying complexity. Frames that fall in well-populated regions under many representations score highly, whereas frames that are outliers under most of them score low.

The score is built from Gaussian mixture models (GMMs). For each feature representation and each number of mixture components d, swept from 2 to 8, a GMM was fit to the per-frame features and used to assign every frame a log-likelihood *L_i_*. The log-likelihoods of each model were normalized as *l_i_* =(*L_i_*-*minL*) /*σ_L_*, and the PWCS of a frame was taken as the average of these normalized log-likelihoods over all feature-representation and component-count combinations.

To make the score reflect equilibrium populations rather than raw simulation sampling, each GMM was weighted by the MSM stationary weights. Because the mixture model does not accept per-sample weights directly, it was fit to a resample of the frames drawn with probability proportional to their MSM weight, so that the fitted density approximates the reweighted equilibrium distribution; the normalization *σ_L_* was likewise computed as a weighted standard deviation. Every frame was then scored against this density, so that over or under sampled regions do not distort the result.

In its general form, PWCS operates on low-dimensional representations derived from several feature types. For the TAR^Mut^ analysis reported here, the R-vector and G-vector descriptors were first projected onto low-dimensional spaces using FPCA, SFA and TAE, and PWCS was applied to these projected representations rather than to the raw descriptors. The two register coordinates, CV1 and CV2, are already two-dimensional order parameters, so they entered PWCS directly without further projection.

## Software availability

GitHub: https://github.com/sbhakat/REVEAL

## Acknowledgements

We acknowledge the use of Claude Opus 4.8 in developing the REVEAL software, streamlining data preparation and processing, and assisting with manuscript revisions.

## References

(1) Al-Hashimi, H. M.; Walter, N. G. RNA Dynamics: It Is about Time. Curr. Opin. Struct. Biol. 2008, 18 (3), 321–329. 10.1016/j.sbi.2008.04.004

(2) Ken, M. L.; Roy, R.; Geng, A.; Ganser, L. R.; Manghrani, A.; Cullen, B. R.; Schulze-Gahmen, U.; Herschlag, D.; Al-Hashimi, H. M. RNA Conformational Propensities Determine Cellular Activity. Nature 2023, 617 (7962), 835–841. 10.1038/s41586-023-06080-x

(3) Hall, K. B. Perspective – RNA Dynamics Today and Tomorrow. J. Mol. Biol. 2026, 438 (16), 169857. 10.1016/j.jmb.2026.169857

(4) Major, F. RNA Is a Dancer: Beyond the One-Structure One-Function Paradigm. Biochem. (Lond). 2025, 47 (4), 13–17. 10.1042/bio_2025_135

(5) Wang, W.; Peng, Z.; Yang, J. Predicting RNA 3D Structure and Conformers Using a Pre-Trained Secondary Structure Model and Structure-Aware Attention. Nat. Mach. Intell. 2026, 8 (5), 722–734. 10.1038/s42256-026-01223-x

(6) Watkins, A. M.; Rangan, R.; Das, R. FARFAR2: Improved De Novo Rosetta Prediction of Complex Global RNA Folds. Structure 2020, 28 (8), 963–976.e6. 10.1016/j.str.2020.05.011

(7) Huang, H.; Guan, X.; Li, W.; Zhang, J.; Wang, W. Protein Dynamics Prediction by Integrating Biophysics and Artificial Intelligence. Curr. Opin. Struct. Biol. 2026, 97, 103223. 10.1016/j.sbi.2026.103223

(8) Herron, L.; Qiu, Y.; Verma, A.; Sreyas Adury, V. S.; John, R.; Lee, S.; Mehdi, S.; Sanwal, D.; Schneekloth, J. S.; Tiwary, P. Ab Initio Prediction of RNA Structure Ensembles with RNAnneal. bioRxiv 2026, 2026.02.01.703098. 10.64898/2026.02.01.703098

(9) Wayment-Steele, H. K.; Ojoawo, A.; Otten, R.; Apitz, J. M.; Pitsawong, W.; Hömberger, M.; Ovchinnikov, S.; Colwell, L.; Kern, D. Predicting Multiple Conformations via Sequence Clustering and AlphaFold2. Nature 2024, 625 (7996), 832–839. 10.1038/s41586-023-06832-9

(10) del Alamo, D.; Sala, D.; Mchaourab, H. S.; Meiler, J. Sampling Alternative Conformational States of Transporters and Receptors with AlphaFold2. Elife 2022, 11, e75751. 10.7554/eLife.75751

(11) Lewis, S.; Hempel, T.; Jiménez-Luna, J.; Gastegger, M.; Xie, Y.; Foong, A. Y. K.; García Satorras, V.; Abdin, O.; Veeling, B. S.; Zaporozhets, I.; Chen, Y.; Yang, S.; Foster, A. E.; Schneuing, A.; Nigam, J.; Barbero, F.; Stimper, V.; Campbell, A.; Yim, J.; Lienen, M.; Shi, Y.; Zheng, S.; Schulz, H.; Munir, U.; Sordillo, R.; Tomioka, R.; Clementi, C.; Noé, F. Scalable Emulation of Protein Equilibrium Ensembles with Generative Deep Learning. Science (1979). 2025, 389 (6761), eadv9817. 10.1126/science.adv9817

(12) Jing, B.; Berger, B.; Jaakkola, T. AlphaFold Meets Flow Matching for Generating Protein Ensembles. ArXiv 2024. 10.48550/arXiv.2402.04845

(13) Vani, B. P.; Aranganathan, A.; Wang, D.; Tiwary, P. AlphaFold2-RAVE: From Sequence to Boltzmann Ranking. J. Chem. Theory Comput. 2023, 19 (14), 4351–4354. 10.1021/acs.jctc.3c00290

(14) Gu, X.; Aranganathan, A.; Tiwary, P. Empowering AlphaFold2 for Protein Conformation Selective Drug Discovery with AlphaFold2-RAVE. eLife Sciences Publications, Ltd 2024. 10.7554/elife.99702.2

(15) Meller, A.; Bhakat, S.; Solieva, S.; Bowman, G. R. Accelerating Cryptic Pocket Discovery Using AlphaFold. J. Chem. Theory Comput. 2023, 19 (14), 4355–4363. 10.1021/acs.jctc.2c01189

(16) Bhakat, S.; Strauch, E.-M. Accelerated Sampling of Protein Dynamics Using BioEmu-Augmented Molecular Simulation. J. Chem. Inf. Model. 2026, 66 (12), 7168–7178. 10.1021/acs.jcim.6c01000

(17) Vats, S.; Bobrovs, R.; Söderhjelm, P.; Bhakat, S. AlphaFold-SFA: Accelerated Sampling of Cryptic Pocket Opening, Protein-Ligand Binding and Allostery by AlphaFold, Slow Feature Analysis and Metadynamics. PLoS One 2024, 19 (8), e0307226. 10.1371/journal.pone.0307226

(18) Husic, B. E.; Pande, V. S. Markov State Models: From an Art to a Science. J. Am. Chem. Soc. 2018, 140 (7), 2386–2396. 10.1021/jacs.7b12191

(19) Puglisi, J. D.; Tan, R.; Calnan, B. J.; Frankel, A. D.; Williamson, J. R. Conformation of the TAR RNA-Arginine Complex by NMR Spectroscopy. Science (1979). 1992, 257 (5066), 76–80. 10.1126/science.1621097

(20) Aboul-ela, F.; Karn, J.; Varani, G. Structure of HIV-1 TAR RNA in the Absence of Ligands Reveals a Novel Conformation of the Trinucleotide Bulge. Nucleic Acids Res. 1996, 24 (20), 3974–3981. 10.1093/nar/24.20.3974

(21) Schulze-Gahmen, U.; Echeverria, I.; Stjepanovic, G.; Bai, Y.; Lu, H.; Schneidman-Duhovny, D.; Doudna, J. A.; Zhou, Q.; Sali, A.; Hurley, J. H. Insights into HIV-1 Proviral Transcription from Integrative Structure and Dynamics of the Tat:AFF4:P-TEFb:TAR Complex. Elife 2016, 5, e15910. 10.7554/eLife.15910

(22) Wei, P.; Garber, M. E.; Fang, S.-M.; Fischer, W. H.; Jones, K. A. A Novel CDK9-Associated C-Type Cyclin Interacts Directly with HIV-1 Tat and Mediates Its High-Affinity, Loop-Specific Binding to TAR RNA. Cell 1998, 92 (4), 451–462. 10.1016/S0092-8674(00)80939-3

(23) Dethoff, E. A.; Petzold, K.; Chugh, J.; Casiano-Negroni, A.; Al-Hashimi, H. M. Visualizing Transient Low-Populated Structures of RNA. Nature 2012, 491 (7426), 724–728. 10.1038/nature11498

(24) Lee, J.; Dethoff, E. A.; Al-Hashimi, H. M. Invisible RNA State Dynamically Couples Distant Motifs. Proceedings of the National Academy of Sciences 2014, 111 (26), 9485–9490. 10.1073/pnas.1407969111

(25) Geng, A.; Ganser, L.; Roy, R.; Shi, H.; Pratihar, S.; Case, D. A.; Al-Hashimi, H. M. An RNA Excited Conformational State at Atomic Resolution. Nat. Commun. 2023, 14 (1), 8432. 10.1038/s41467-023-43673-6

(26) Ganser, L. R.; Chu, C.-C.; Bogerd, H. P.; Kelly, M. L.; Cullen, B. R.; Al-Hashimi, H. M. Probing RNA Conformational Equilibria within the Functional Cellular Context. Cell Rep. 2020, 30 (8), 2472–2480.e4. 10.1016/j.celrep.2020.02.004

(27) Ganser, L. R.; Kelly, M. L.; Patwardhan, N. N.; Hargrove, A. E.; Al-Hashimi, H. M. Demonstration That Small Molecules Can Bind and Stabilize Low-Abundance Short-Lived RNA Excited Conformational States. J. Mol. Biol. 2020, 432 (4), 1297–1304. 10.1016/j.jmb.2019.12.009

(28) Watkins, A. M.; Rangan, R.; Das, R. FARFAR2: Improved De Novo Rosetta Prediction of Complex Global RNA Folds. Structure 2020, 28 (8), 963–976.e6. 10.1016/j.str.2020.05.011

(29) Roy, R.; Geng, A.; Shi, H.; Merriman, D. K.; Dethoff, E. A.; Salmon, L.; Al-Hashimi, H. M. Kinetic Resolution of the Atomic 3D Structures Formed by Ground and Excited Conformational States in an RNA Dynamic Ensemble. J. Am. Chem. Soc. 2023, 145 (42), 22964–22978. 10.1021/jacs.3c04614

(30) Röder, K. Decoding RNA Structural Ensembles: Energy Landscape Exploration of the TAR Stemloop. J. Chem. Theory Comput. 2026, 22 (2), 1111–1121. 10.1021/acs.jctc.5c01677

(31) Abramson, J.; Adler, J.; Dunger, J.; Evans, R.; Green, T.; Pritzel, A.; Ronneberger, O.; Willmore, L.; Ballard, A. J.; Bambrick, J.; Bodenstein, S. W.; Evans, D. A.; Hung, C.-C.; O’Neill, M.; Reiman, D.; Tunyasuvunakool, K.; Wu, Z.; Žemgulytė, A.; Arvaniti, E.; Beattie, C.; Bertolli, O.; Bridgland, A.; Cherepanov, A.; Congreve, M.; Cowen-Rivers, A. I.; Cowie, A.; Figurnov, M.; Fuchs, F. B.; Gladman, H.; Jain, R.; Khan, Y. A.; Low, C. M. R.; Perlin, K.; Potapenko, A.; Savy, P.; Singh, S.; Stecula, A.; Thillaisundaram, A.; Tong, C.; Yakneen, S.; Zhong, E. D.; Zielinski, M.; Žídek, A.; Bapst, V.; Kohli, P.; Jaderberg, M.; Hassabis, D.; Jumper, J. M. Accurate Structure Prediction of Biomolecular Interactions with AlphaFold 3. Nature 2024, 630 (8016), 493–500. 10.1038/s41586-024-07487-w

(32) Candido, S.; Hayes, T.; Derry, A.; Rao, R.; Lin, Z.; Verkuil, R.; Wu, B. Z.; Lee, J. S.; Bruguera, E. S.; Keval, J. A.; Kopylov, M.; Pak, J. E.; Wu, W.; Thomas, N.; Mataraso, S.; Hsu, A.; Trotman-Grant, A. C.; Fatras, K.; dos Santos Costa, A.; Badkundri, R.; Akın, H.; Oktay, D.; Deaton, J.; Montabana, E.; Sitwala, H.; Yu, Y.; Wiggert, M.; Carlin, D. A.; Goering, A. W.; Blazejewski, T.; Sandora, M.; Hla, M.; Jia, T. Z.; Kloker, L. H.; Sofroniew, N. J.; Uehara, M.; Pannu, J.; Bachas, S.; Liu, D. S.; Sercu, T.; Rives, A. Language Modeling Materializes a World Model of Protein Biology. bioRxiv 2026, 2026.06.03.729735. 10.64898/2026.06.03.729735

(33) The OpenFold3 Team. OpenFold3-Preview. 2025.

(34) Passaro, S.; Corso, G.; Wohlwend, J.; Reveiz, M.; Thaler, S.; Somnath, V. R.; Getz, N.; Portnoi, T.; Roy, J.; Stark, H.; Kwabi-Addo, D.; Beaini, D.; Jaakkola, T.; Barzilay, R. Boltz-2: Towards Accurate and Efficient Binding Affinity Prediction. bioRxiv 2025, 2025.06.14.659707. 10.1101/2025.06.14.659707

(35) Zhang, Y.; Gong, C.; Sun, J.; Guan, J.; Ren, M.; Xue, S.; Zhang, H.; Ma, W.; Liu, Z.; Chen, X.; Xiao, W. Protenix-v2: Broadening the Reach of Structure Prediction and Biomolecular Design. bioRxiv 2026, 2026.04.10.717613. 10.64898/2026.04.10.717613

(36) Lorenz, R.; Bernhart, S. H.; Höner zu Siederdissen, C.; Tafer, H.; Flamm, C.; Stadler, P. F.; Hofacker, I. L. ViennaRNA Package 2.0. Algorithms for Molecular Biology 2011, 6 (1), 26. 10.1186/1748-7188-6-26

(37) Zhang, H.; Li, S.; Zhang, L.; Mathews, D. H.; Huang, L. LazySampling and LinearSampling: Fast Stochastic Sampling of RNA Secondary Structure with Applications to SARS-CoV-2. Nucleic Acids Res. 2023, 51 (2), e7–e7. 10.1093/nar/gkac1029

(38) Ding, Y.; Chan, C. Y.; Lawrence, C. E. Sfold Web Server for Statistical Folding and Rational Design of Nucleic Acids. Nucleic Acids Res. 2004, 32 (suppl_2), W135–W141. 10.1093/nar/gkh449

(39) Bhakat, S. Collective Variable Discovery in the Age of Machine Learning: Reality, Hype and Everything in Between. RSC Adv. 2022, 12 (38), 25010–25024. 10.1039/D2RA03660F

(40) Roy, R.; Geng, A.; Shi, H.; Merriman, D. K.; Dethoff, E. A.; Salmon, L.; Al-Hashimi, H. M. Kinetic Resolution of the Atomic 3D Structures Formed by Ground and Excited Conformational States in an RNA Dynamic Ensemble. J. Am. Chem. Soc. 2023, 145 (42), 22964–22978. 10.1021/jacs.3c04614

(41) Röblitz, S.; Weber, M. Fuzzy Spectral Clustering by PCCA+: Application to Markov State Models and Data Classification. Adv. Data Anal. Classif. 2013, 7 (2), 147–179. 10.1007/s11634-013-0134-6

(42) Mardt, A.; Pasquali, L.; Wu, H.; Noé, F. VAMPnets for Deep Learning of Molecular Kinetics. Nat. Commun. 2018, 9 (1), 5. 10.1038/s41467-017-02388-1

(43) Bhakat, S.; Vats, S.; Mardt, A.; Degterev, A. Generalizable Protein Dynamics in Kinases: Physics Is the Key. bioRxiv 2025, 2025.03.06.641878. 10.1101/2025.03.06.641878

(44) Bonati, L.; Piccini, G.; Parrinello, M. Deep Learning the Slow Modes for Rare Events Sampling. Proceedings of the National Academy of Sciences 2021, 118 (44), e2113533118. 10.1073/pnas.2113533118

(45) Viegas, R. G.; Sinha, A.; Mainan, A.; Sanbonmatsu, K. Y.; Onuchic, J. N.; Roy, S.; Leite, V. B. P. Critical Role of Mg2+ Ions in RNA Folding Transitions: Anchoring the A-Minor Twist in the SAM-II Riboswitch. J. Phys. Chem. B 2025, 129 (36), 9058–9067. 10.1021/acs.jpcb.5c02586

(46) Fischer, N. M.; Polêto, M. D.; Steuer, J.; van der Spoel, D. Influence of Na+ and Mg2+ Ions on RNA Structures Studied with Molecular Dynamics Simulations. Nucleic Acids Res. 2018, 46 (10), 4872–4882. 10.1093/nar/gky221

(47) Choi, T.; Li, Z.; Song, G.; Chen, H.-F. Comprehensive Comparison and Critical Assessment of RNA-Specific Force Fields. J. Chem. Theory Comput. 2024, 20 (6), 2676–2688. 10.1021/acs.jctc.4c00066

(48) Mlýnský, V.; Kührová, P.; Pykal, M.; Krepl, M.; Stadlbauer, P.; Otyepka, M.; Banáš, P.; Šponer, J. Can We Ever Develop an Ideal RNA Force Field? Lessons Learned from Simulations of the UUCG RNA Tetraloop and Other Systems. J. Chem. Theory Comput. 2025, 21 (8), 4183–4202. 10.1021/acs.jctc.4c01357

(49) Singh, O.; Venugopal, P. P.; Chakraborty, D. Effect of Water Models on The Stability of RNA: Role of Counter-Ions. Chemical Physics Impact 2023, 7, 100313. 10.1016/j.chphi.2023.100313

(50) Linzer, J. T.; Aminov, E.; Abdullah, A. S.; Kirkup, C. E.; Diaz Ventura, R. I.; Bijoor, V. R.; Jung, J.; Huang, S.; Tse, C. G.; Álvarez Toucet, E.; Onghai, H. P.; Ghosh, A. P.; Grodzki, A. C.; Haines, E. R.; Iyer, A. S.; Khalil, M. K.; Leong, A. P.; Neuhaus, M. A.; Park, J.; Shahid, A.; Xie, M.; Ziembicki, J. M.; Simmerling, C.; Nagan, M. C. Accurately Modeling RNA Stem-Loops in an Implicit Solvent Environment. J. Chem. Inf. Model. 2024, 64 (15), 6092–6104. 10.1021/acs.jcim.4c00756

(51) Krepl, M.; Mlýnský, V.; Rusnáková, A.; Banáš, P.; Otyepka, M.; Šponer, J. Destabilization of Structured RNAs by OPC and TIP4PD Water Models. J. Chem. Theory Comput. 2026, 22 (4), 2013–2027. 10.1021/acs.jctc.5c01678

(52) Haller, A.; Soulière, M. F.; Micura, R. The Dynamic Nature of RNA as Key to Understanding Riboswitch Mechanisms. Acc. Chem. Res. 2011, 44 (12), 1339–1348. 10.1021/ar200035g

(53) Montange, R. K.; Batey, R. T. Riboswitches: Emerging Themes in RNA Structure and Function. Annu. Rev. Biophys. 2008, 37 (Volume 37, 2008), 117–133. 10.1146/annurev.biophys.37.032807.130000

(54) Townley, J.; Kladwang, W.; Baker, D.; Blair, H. M.; Choe, C. A.; El Nesr, G.; Favor, A.; Fisker, E.; Haack, D. B.; He, S.; Hingey, J.; Huang, P.-S.; Huang, R.; Joshi, C. K.; Karagianes, T.; Kubaney, A.; Liò, P.; Mancino, A.; Romano, J.; Rudolfs, B.; Spellmon, N.; Toor, N.; Verma, J.; Wu, V.; Yu, Z.; Participants, E.; Das, R. De Novo Design of RNA Pseudoknots with Deep Learning. bioRxiv 2026, 2026.05.21.726960. 10.64898/2026.05.21.726960

(55) Viegas, R. G.; Sinha, A.; Mainan, A.; Sanbonmatsu, K. Y.; Onuchic, J. N.; Roy, S.; Leite, V. B. P. Critical Role of Mg2+ Ions in RNA Folding Transitions: Anchoring the A-Minor Twist in the SAM-II Riboswitch. J. Phys. Chem. B 2025, 129 (36), 9058–9067. 10.1021/acs.jpcb.5c02586

(56) Zgarbová, M.; Otyepka, M.; Šponer, J.; Mládek, A.; Banáš, P.; Cheatham III, T. E.; Jurečka, P. Refinement of the Cornell et al. Nucleic Acids Force Field Based on Reference Quantum Chemical Calculations of Glycosidic Torsion Profiles. J. Chem. Theory Comput. 2011, 7 (9), 2886–2902. 10.1021/ct200162x

(57) Jorgensen, W. L.; Chandrasekhar, J.; Madura, J. D.; Impey, R. W.; Klein, M. L. Comparison of Simple Potential Functions for Simulating Liquid Water. J. Chem. Phys. 1983, 79 (2), 926–935. 10.1063/1.445869

(58) Abraham, M. J.; Murtola, T.; Schulz, R.; Páll, S.; Smith, J. C.; Hess, B.; Lindahl, E. GROMACS: High Performance Molecular Simulations through Multi-Level Parallelism from Laptops to Supercomputers. SoftwareX 2015, 1–2, 19–25. 10.1016/j.softx.2015.06.001

(59) Parrinello, M.; Rahman, A. Polymorphic Transitions in Single Crystals: A New Molecular Dynamics Method. J. Appl. Phys. 1981, 52 (12), 7182–7190. 10.1063/1.328693

(60) Darden, T.; York, D.; Pedersen, L. Particle Mesh Ewald: An N⋅log(N) Method for Ewald Sums in Large Systems. J. Chem. Phys. 1993, 98 (12), 10089–10092. 10.1063/1.464397

(61) Hess, B.; Bekker, H.; Berendsen, H. J. C.; Fraaije, J. G. E. M. LINCS: A Linear Constraint Solver for Molecular Simulations. J. Comput. Chem. 1997, 18 (12), 1463–1472. 10.1002/(SICI)1096-987X(199709)18:12<1463::AIDJCC4>3.0.CO;2-H

(62) McGibbon, R. T.; Beauchamp, K. A.; Harrigan, M. P.; Klein, C.; Swails, J. M.; Hernández, C. X.; Schwantes, C. R.; Wang, L.-P.; Lane, T. J.; Pande, V. S. MDTraj: A Modern Open Library for the Analysis of Molecular Dynamics Trajectories. Biophys. J. 2015, 109 (8), 1528–1532. 10.1016/j.bpj.2015.08.015

(63) Bottaro, S.; Di Palma, F.; Bussi, G. The Role of Nucleobase Interactions in RNA Structure and Dynamics. Nucleic Acids Res. 2014, 42 (21), 13306–13314. 10.1093/nar/gku972

(64) Konovalov, K. A.; Unarta, I. C.; Cao, S.; Goonetilleke, E. C.; Huang, X. Markov State Models to Study the Functional Dynamics of Proteins in the Wake of Machine Learning. JACS Au 2021, 1 (9), 1330–1341. 10.1021/jacsau.1c00254

(65) Pande, V. S.; Beauchamp, K.; Bowman, G. R. Everything You Wanted to Know about Markov State Models but Were Afraid to Ask. Methods 2010, 52 (1), 99–105. 10.1016/j.ymeth.2010.06.002

(66) Suárez, E.; P. Wiewiora, R.; Wehmeyer, C.; Noé, F.; D. Chodera, J.; M. Zuckerman, D. What Markov State Models Can and Cannot Do: Correlation versus Path-Based Observables in Protein-Folding Models. J. Chem. Theory Comput. 2021, 17 (5), 3119–3133. 10.1021/acs.jctc.0c01154

(67) Scherer, M. K.; Trendelkamp-Schroer, B.; Paul, F.; Pérez-Hernández, G.; Hoffmann, M.; Plattner, N.; Wehmeyer, C.; Prinz, J.-H.; Noé, F. PyEMMA 2: A Software Package for Estimation, Validation, and Analysis of Markov Models. J. Chem. Theory Comput. 2015, 11 (11), 5525–5542. 10.1021/acs.jctc.5b00743

