## Supporting Information for "REVEAL: RNA Ensemble Validation and Exploration with AI-guided Landscapes"

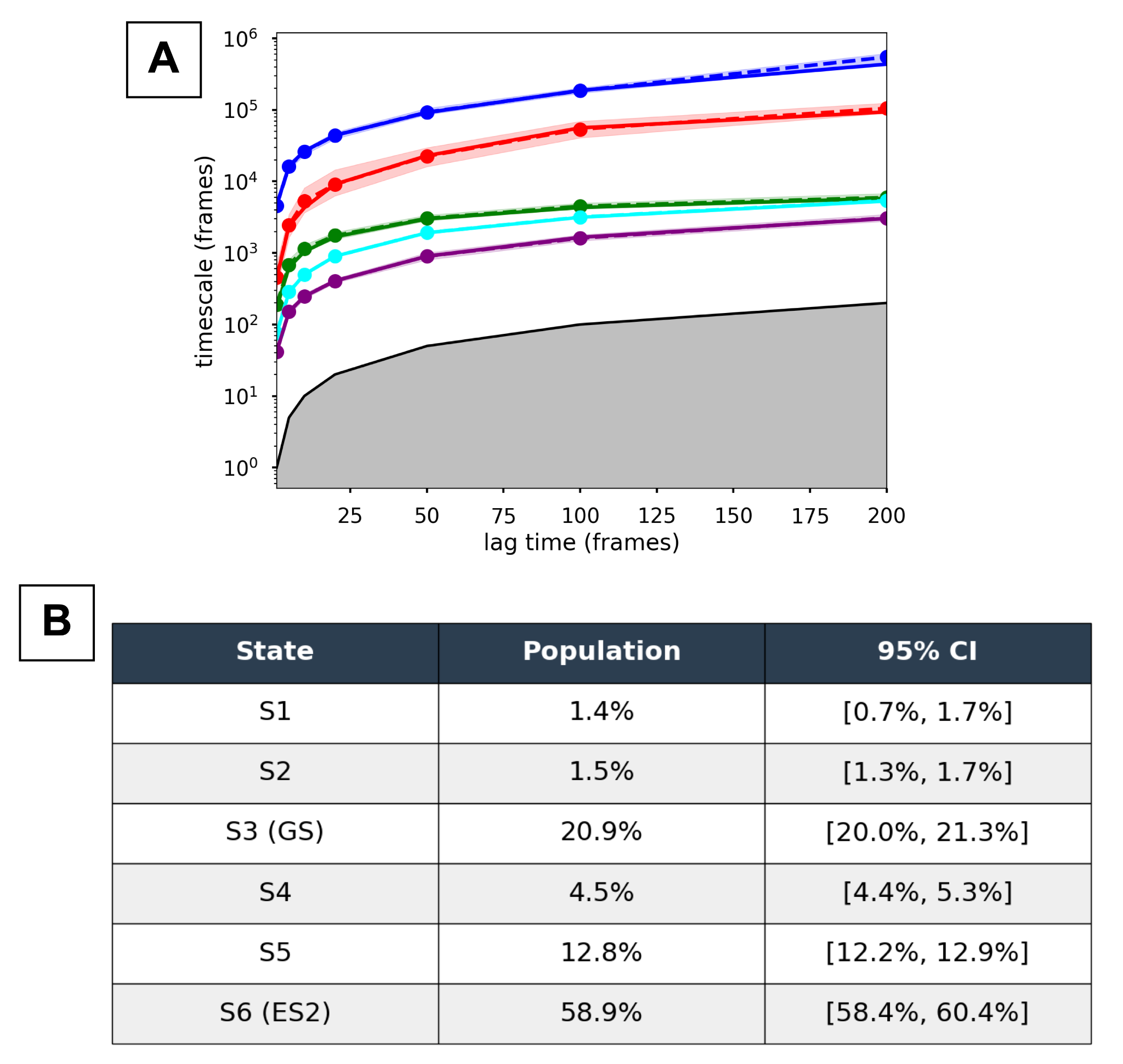

Figure S1. Implied timescales and metastable state populations. (A) Implied timescales (in frames) of the Markov state model as a function of lag time; each colored line is a resolved event with its 95% confidence interval (shaded), and the gray region marks timescales faster than the lag time that cannot be resolved. (B) Equilibrium populations of the six metastable states (S1–S6) with 95% confidence intervals; S3 is the ground state (GS) and S6 the dominant excited state (ES2).

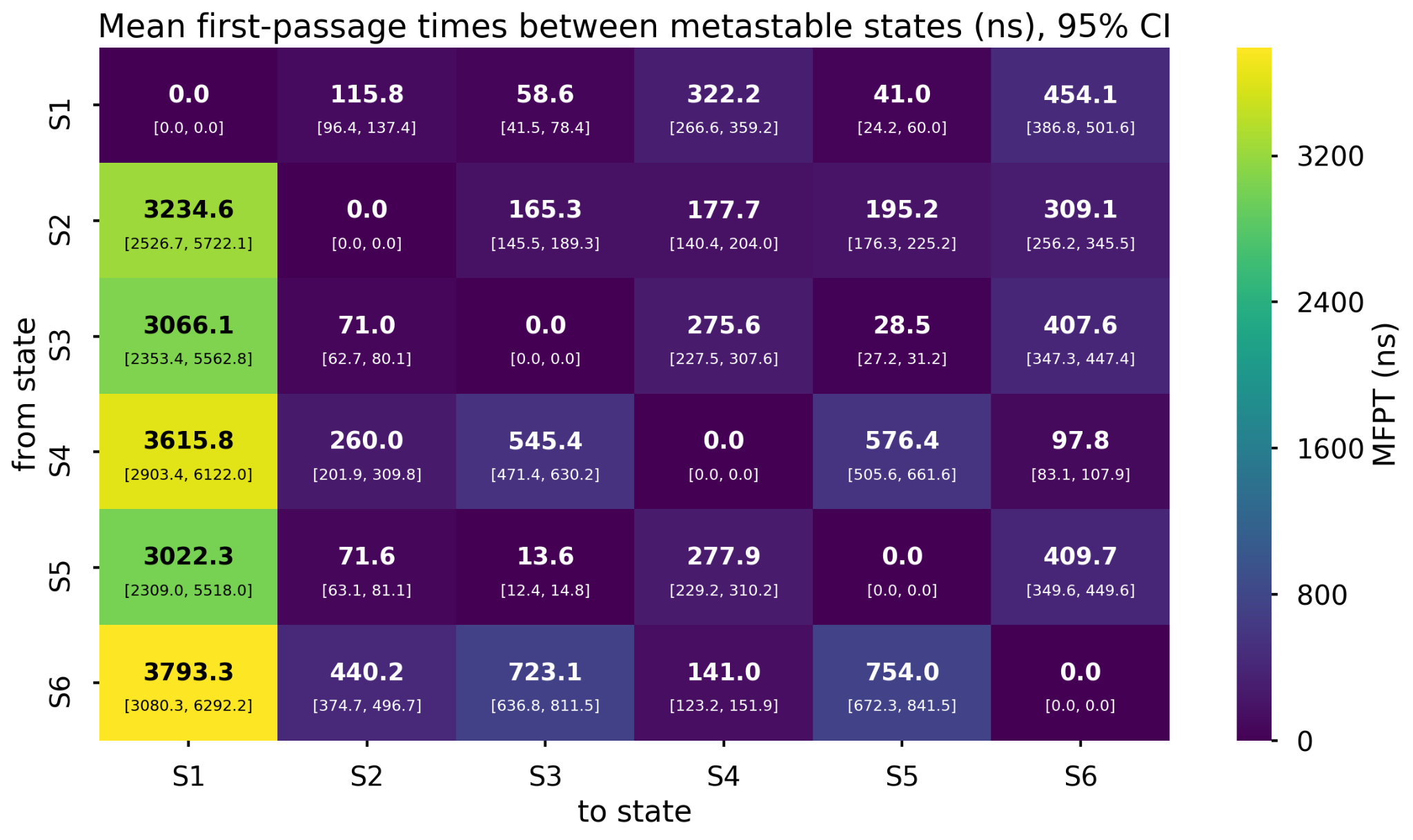

Figure S2. Mean first-passage times (MFPT) between metastable states. Heatmap showing MFPT (in ns) to travel from each starting state (rows, S1–S6) to each target state (columns, S1–S6). Values in brackets are 95% confidence intervals, and color indicates MFPT magnitude.

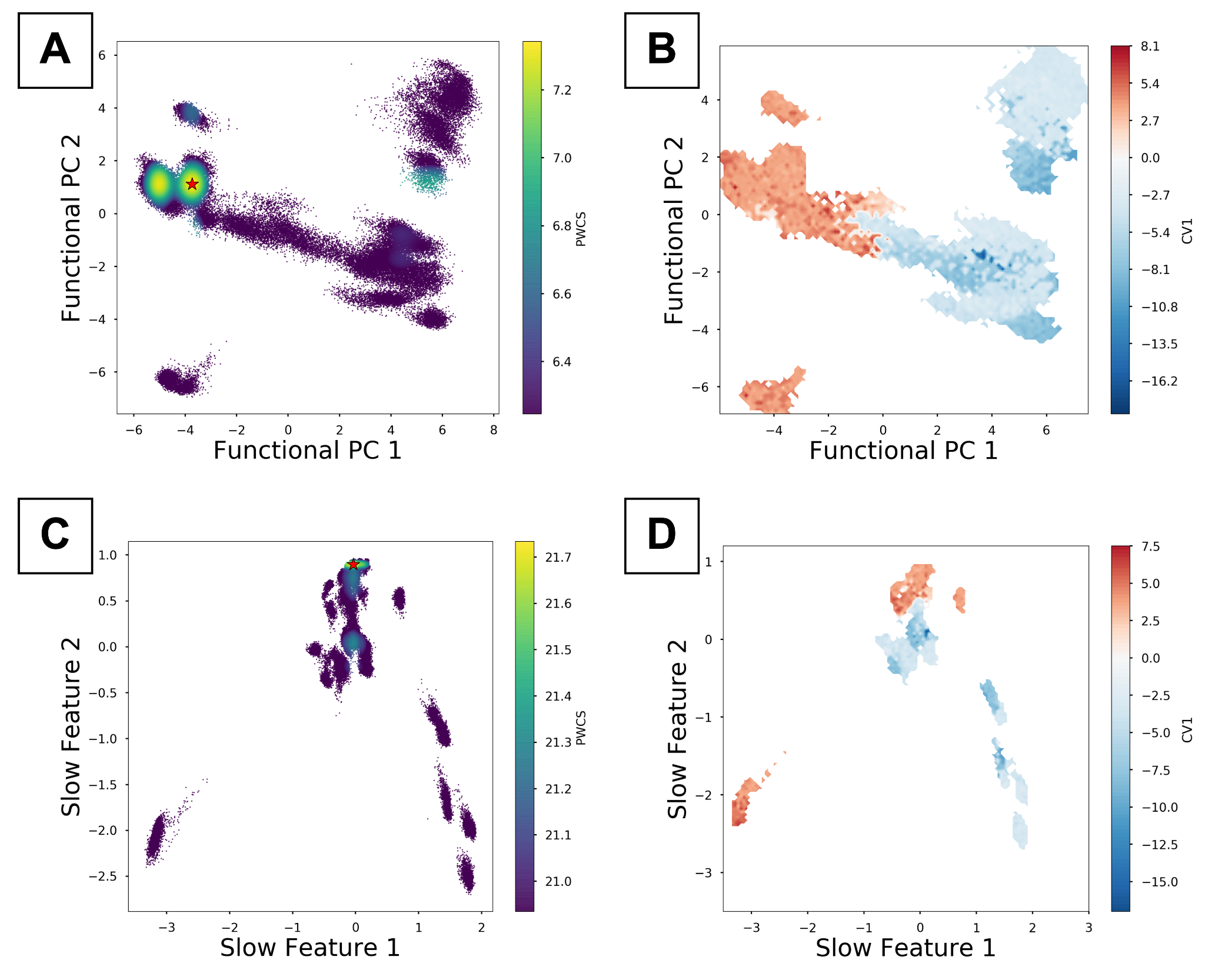

Figure S3. Conformational landscapes projected onto reduced coordinate spaces. (A, B) Functional principal components (PC 1 vs PC 2) colored by PWCS (A) and by CV1 (B). (C, D) Slow features (Slow Feature 1 vs 2) colored by PWCS (C) and by CV1 (D). The red star marks the global free energy minimum which corresponds to the dominant ES2 state of TARMut. Positive values of CV1 belong to ES2 like states.

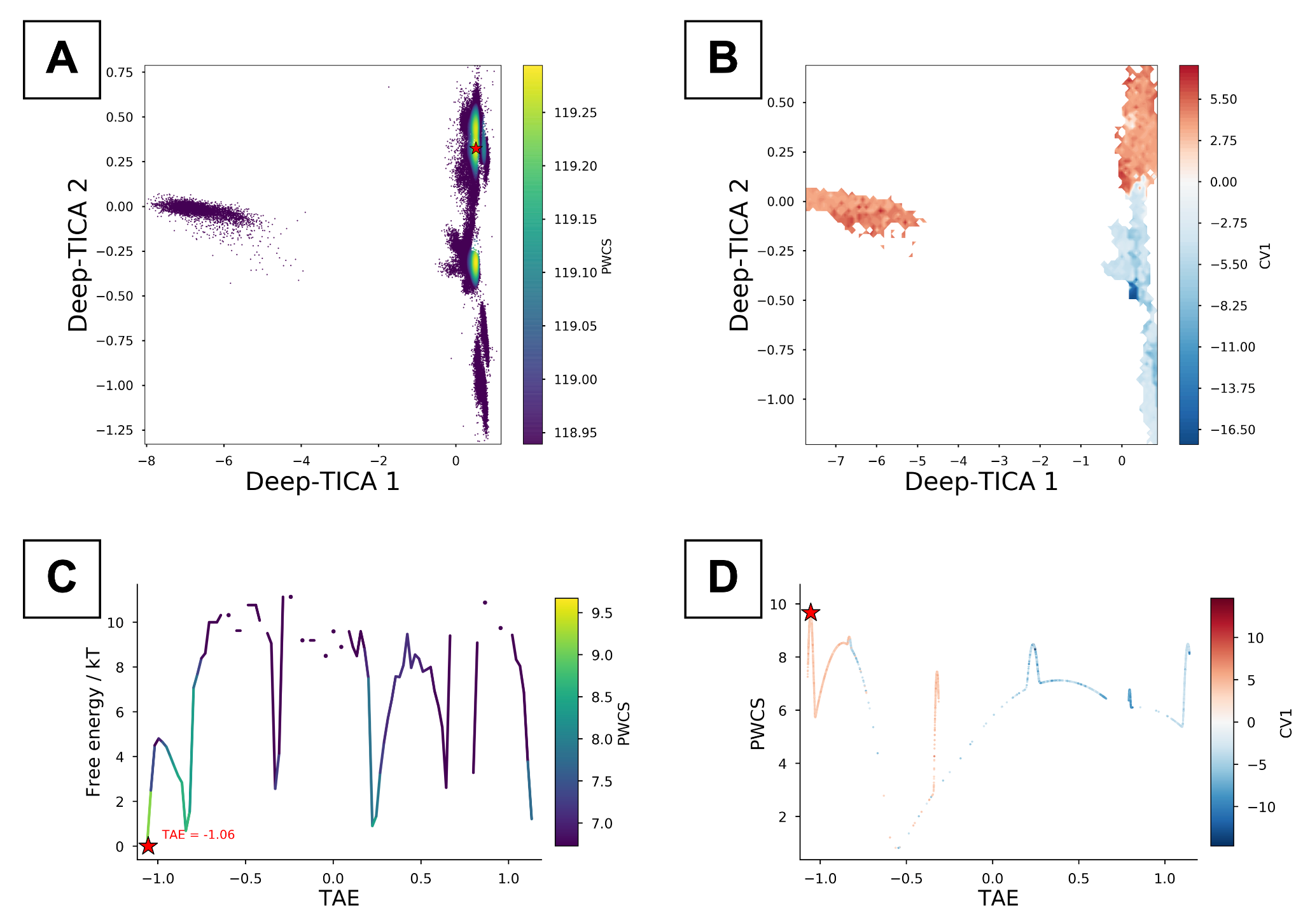

Figure S4. Deep-TICA landscape and free energy along the TAE coordinate. (A, B) Deep-TICA components (1 vs 2) colored by PWCS (A) and by CV1 (B). (C) Free energy (in kT) along the TAE coordinate colored by PWCS, with the minimum at TAE = −1.06 marked (red star). (D) PWCS versus TAE colored by CV1; the red star marks the global free energy minimum corresponding to the dominant ES2 state. Positive values of CV1 belong to ES2 like states.
